# Beyond Metabolism: Pyruvate Carboxylase Acts as a Sequence-Selective Small RNA Sensor Orchestrating Antiviral Responses

**DOI:** 10.64898/2026.06.29.735367

**Authors:** Udeshika Kariyawasam, Suranjana Goswami, Ming Hao, Rosana Wiscovitch-Russo, Qian Chen, Jun Yang, Ju Qu, Mayra Marquez, Hongyan Sui, Weizhong Chang, Tomozumi Imamichi

## Abstract

Interleukin-27 (IL-27) is an anti-HIV cytokine that induces 14 novel microRNAs (miRNAs) in primary CD4(+) T cells. We previously reported that transfection of two of these miRNA mimics, miRTC10 and miRTC14 into human primary macrophages, differentially induced interferon (IFN)-α2, -α8, -α13, and -λ1 expression. However, the mechanism underlying this activation remains unclear. Here, we demonstrate that miRTC14 does not directly target canonical IFN-regulatory genes but instead engages with pyruvate carboxylase (PC) and laboratory of genetics and physiology-2 (LGP2/DHX58) as direct binding partner proteins. Functional analysis revealed that miRTC14 transfection induces IFN expression by more than 100-fold (p<0.001), whereas PC or LGP2 depletion by siRNAs markedly attenuated this response (50–100-fold reduction, p<0.01). Reconstitution of PC and LGP2 in deficient HEK293 cells restored miRTC14-driven IFN induction. Notably, IFN activation depended on sequence features at the duplex termini. However, PC–miRTC14–LGP2 axis activates TBK1-dependent phosphorylation of IRF3/7, similarly to canonical RNA sensors (RIG-I/MDA5), but this process induced differential IFN subtype. These findings establish PC as a miRNA-binding protein and define a previously unrecognized RNA-sensing mechanism, linking metabolic enzymes to RNA sequence-dependent innate immunity.

**GRAPHICAL ABSTRACT:** Graphical Abstract:
microRNA sequence–dependent activation of innate immunity by pyruvate carboxylase (PC).
PC–miRTC14–LGP2 axis induces TBK1 phosphorylation via an as-yet-unidentified adapter(s). The activated TBK1 (p-TBK1) subsequently phosphorylates IRF3 and/or IRF7 leading to selective IFN subtypes (IFN-α2, IFN-α8, IFN-α13, and IFN-λ1). (Created in BioRender. Kariyawasam, U. (2027) https://BioRender.com/u82s04p)

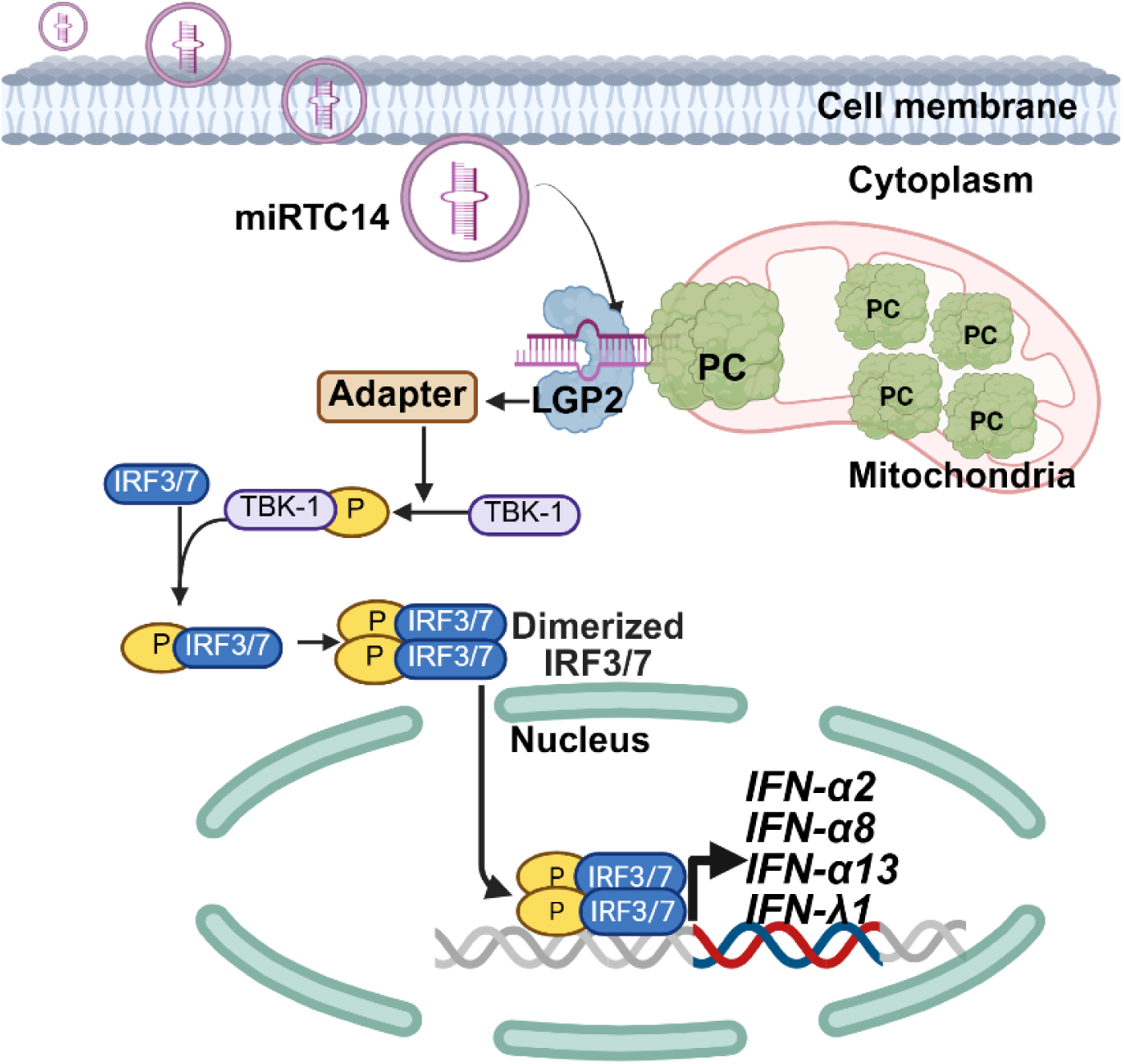

## INTRODUCTION

Pyruvate carboxylase (PC) is a member of the biotin-dependent carboxylase family and is widely distributed among both prokaryotic and eukaryotic organisms (1–4). In eukaryotic cells, PC is predominantly localized to the mitochondria, where it plays essential roles in gluconeogenesis and lipogenesis. PC catalyses the ATP-dependent carboxylation of pyruvate to generate oxaloacetate (OAA). During gluconeogenesis, OAA is converted to phosphoenolpyruvate and subsequently utilized for glucose synthesis. During lipogenesis, OAA is converted to citrate, which is transported to cytoplasm and serves as a source of acetyl-CoA for fatty acid synthesis. Through its anaplerotic function, PC replenishes the tricarboxylic acid cycle (TCA) cycle intermediates that are consumed during biosynthetic processes, thereby maintaining cellular metabolic homeostasis. Many cancer cells exploit this metabolic function to support proliferation, survival, and adaptation to metabolic stress. Consequently, PC has been implicated in tumour progression and cancer cell metabolism (4,5). Beyond its canonical metabolic functions, PC has non-enzymatic function as a moonlighting protein (3,6,7). PC has been reported to possess anti-senescence activity through the regulation of the tumour suppressor p53 pathway. In addition, recent studies have linked PC to antiviral responses and the suppression of viral replication (8–10) and been associated with the induction of interferon (IFN) production (6). These findings suggest that PC performs functions beyond metabolism and may contribute to the regulation of innate immune responses. However, whether PC directly participates in the recognition of immunostimulatory RNAs and the initiation of antiviral IFN responses remains unknown.

The innate immune system provides the first line of defence against microbial invasion. Pathogen-associated molecules are detected by pattern-recognition receptors (PRRs), including Toll-like receptors (TLRs), RIG-I-like receptors (RLRs), DNA sensors (11), and the initiate innate immune responses. RLRs are a family of three cytosolic RNA recognition receptors consisting of RIG-I, melanoma differentiation-associated gene-5 (MDA5), and laboratory of genetics and physiology-2 (LGP2) (11). These proteins play key roles for detecting viral RNA in the cytoplasm and initiating innate immune responses (12). Upon detecting RNA, RIG-I and MDA5 activate downstream signalling through TANK-binding kinase-1 (TBK1) to induce IFNs (13). LGP2 is expressed at low basal levels but is upregulated during infection and can bind diverse RNA species independent of length or 5′-phosphate status (14–16). Although LGP2 lacks canonical signalling domains (12) and was initially considered a negative regulator (15), genetic studies have established LGP2 as a positive modulator of RIG-I and MDA5 signalling (12). Although significant progress has been made, current understanding of RNA sensing is primarily based on structural recognition, leaving the potential for sequence-specific intracellular RNA sensors largely unexplored.

microRNAs (miRNAs) are ∼19–24 nucleotide non-coding small RNAs that regulate gene expression post-transcriptionally through sequence-specific interactions with target mRNAs (17). miRNAs are generated from precursor transcripts through sequential processing by RNase III enzymes, including Dicer, resulting in the formation of a small RNA duplex composed of a guide strand and a passenger strand (17). Strand selection is largely determined by thermodynamic asymmetry, favouring incorporation of the guide strand into Argonaute (Ago) complexes, while the passenger strand is typically degraded (17,18). However, accumulating evidence suggests that passenger strand can also contribute to the regulation of miRNA homeostasis and downstream gene regulatory networks (19,20). Synthetic mature miRNA (miRNA mimics) are widely used to study their functions and for therapeutic applications (21,22). miRNA mimics also form duplexes as do endogenous miRNAs. However, emerging evidence suggests that certain miRNA mimics can exert non-canonical functions, including off target innate immune activation or endosomal stresses (18,23,24). As part of innate immune responses, transfected synthetic small RNA has been reported to induce type I and Type III IFNs (24). Therefore, certain off-target effects reflect engagement of innate immune responses. However, the mechanisms underlying such small RNA-induced IFN responses remain poorly defined.

We previously identified three novel anti-HIV miRNAs (miRTC10, miRTC14 and miRAB40) in HIV-resist human primary T cells and macrophages differentiated or polarized by anti-HIV cytokine, interleukin-27 (IL-27) (25,26). In the current study, we compared anti-HIV effect among the miRNAs and demonstrate that a specific anti-HIV miRNA induces robust and selective expression of IFN-α2, IFN-α8, IFN-α13 and IFN-λ1 in human macrophages. We demonstrate that mechanistically, this response is mediated through an unexpected pathway involving PC and the LGP2–TBK1 signalling axis. Strikingly, we identified PC, as a previously unrecognized RNA sensor that recognizes RNA in a sequence-dependent manner. These findings reveal a non-canonical mechanism of innate immune activation that links cellular metabolism to RNA sensing and establish a conceptual framework in which metabolic enzymes function as regulators of antiviral immunity. This work provides new insight into RNA-driven immune activation and highlights the potential of miRNA-based strategies to selectively engage antiviral-IFN responses against HIV and IFN susceptible pathogens, and our data extends the moonlighting function of PC.

## MATERIAL AND METHODS

### Cells and reagents

PBMCs were isolated from healthy donors’ apheresis packs from the National Institutes of Health Blood Bank (Bethesda, MD, USA) or STEM CELL (Cambridge, MA, USA) using a lymphocyte separation medium (ICN Biomedical, Aurora, OH, USA) (27). CD14(+) monocytes were separated from PBMCs using CD14 MicroBeads (Miltenyi Biotec, Auburn, CA, USA), according to the manufacturer’s instructions. The purity of the cell types was at least 90%, based on the flow cytometric analysis. Cell viability was determined using the Trypan blue (Thermo Fisher Scientific, Waltham, MA, USA) exclusion method. The CD14(+) monocyte-derived macrophages (MDMs) were differentiated from CD14(+) monocytes using 25 ng/mL of macrophage colony–stimulating factor (M-CSF) (R&D Systems, Minneapolis, MN, USA) in macrophage serum-free media (M-SFM, Thermo Fisher Scientific) as previously described. Briefly, 10×10^6^ monocytes were cultured in 10 mL of M-SFM, supplemented with 10 μg/mL Gentamicin (Thermo Fisher Scientific, Cat# 15710-072) and 10 mM 4-(2-hydroxyethyl)-1-piperazine ethane sulfonic acid (HEPES), pH 7.3 (Quality Biological, Gaithersburg, MD, USA, cat # 118-089-721) in the presence of 25 ng/mL of M-CSF in a 100 mm Petri dish at 37°C and 5% CO_2_ with saturating humidity for 7-days. Half of medium was replaced with fresh M-SFM on the 4th day. After differentiation, MDMs were maintained in the complete D-MEM medium (Thermo Fisher) supplemented with 10% (v/v) fetal bovine serum (FBS; R&D Systems), 10 mM HEPES, and 10 μg/mL Gentamicin (D10 medium) as previously described (27,28). HEK293T cells were obtained from ATCC (Manassas, VA, USA) and maintained in D10 medium. HEK293-PC-KO cell line and HEK293-LGP2-KO cell line were purchased from Ubigene (Austin, TX, USA) and maintained in D10 medium.

### Preparation of HIVAD8 replication competent virus and infection assay

To assess the impact of each miRNA on HIV replication in MDMs, HIVAD8 virus was used. HIVAD8 was prepared by transfection of plasmid encoding full length of HIV_AD8_ strain into 293T cells (29). This process was performed using Transit 293 (Mirus, Madison, WI) as previously described (28). The infectious titter (TCID50/ml) of the virus stock was determined by an endpoint assay as described before(25). MDMs (50×10^3^ cells / 96-well) that had been transfected with miRNA were infected at 5000 TCID50/10^6^ cells for 2 hr at 37 °C. Then the cells were cultured for 14 days with half the media being changed every 3–4 days with fresh D10, HIV replication was quantified using p24 antigen capture kit (Revvity, Waltham, MA, USA) using a SpectraMax M5 Multi-Mode Microplate Reader (Molecular Devices, San Jose, California).

### Preparation of HIVLuc-VSVG and infection

To assess the impact of each miRNA on HIV infection, replication incompetent pseudo-typed HIVLuc-VSVG virus was used (27,30). The pseudo typed HIV was produced by co-transfecting 293T cells with pNL4-3Luc (8 μg) and pLTR-VSVG (4 μg) using a TransIT-293 transfection kit (Mirus, Madison, WI). Virus containing supernatants were harvested 48 h after transfection and filtered with 0.45μm Steriflip Filter Units (MilliporeSigma, Burlington, MA, USA). Virus particles were pelleted by ultracentrifugation (100,000 x g at 4^0^C) through a 20% sucrose-10 mM HEPES (pH7.3) cushion and resuspended in PBS. Virus amounts were quantified using p24 antigen kit (27). MDMs (50×10^3^ cell/well) were seeded in 96-well plated and cultured for 16 hr. The cells were then transfected with 10 nM of miRNAs for 48 hr and then infected with 100 ng p24/ml of HIVLuc-VSVG for 2 hr at 37°C. After infection, HIVLuc–VSVG-infected cells were washed by pipetting and then cultured for two days in 200 μL D10. All assays were conducted quadruplicated. The Infected cell amounts were quantified using the Bright-Glo Luciferase kit (Promega, Madison, WI) with a TECAN SPARK plate reader (TECAN, Männedorf, Switzerland).

### B18R experiment

After the cells were transfected with miRNA, the cells were treated with and without B18R (1 μg/ml, recombinant vaccine virus-encoded IFN inhibitor) (R&D Systems). For control purpose a set of MDMs were just treated with IFNα (100 U/ml) (R&D Systems). It was kept for 72hrs, and the cells were infected with HIVLUC-VSVG for 48 hrs and the percent replication was measured.

### Gel electrophoresis and western blot

Western blotting (WB) was performed as previously described (31). Briefly, control or transfected cells in 6-well plates (1.5×10^6^ cells/well) were washed three times with cold PBS, lysed using 150 μL of 1x Radioimmunoprecipitation assay (RIPA) lysis buffer (Boston BioProducts, Milford, MA, USA) supplemented with 5 mM EDTA (Thermo Fisher) and 1x phosphate and protease inhibitor cocktail (Thermo Fisher Scientific), and incubated on ice for 15 min. Then, the cell debris-free cell lysates were obtained by centrifuging at 15,000×g at 4 °C for 10 min. Protein concentration was determined using the BCA protein assay kit (Pierce, Thermo Fisher Scientific), and 20 μg total protein for each lysate was subjected to 4%–12% NuPAGE Bis-Tris gels (Thermo Fisher Scientific) in MOPS buffer (Thermo Fisher Scientific) under reducing condition. Proteins were transferred onto 0.45-μm pore-size nitrocellulose membranes (Thermo Fisher Scientific) and probed with antibodies, anti-PC Rabbit Ab (Cat# 49381),, anti-p-TBK1 (S172) (D52C2) XP(R) Rabbit mAb (Cat# 5483), anti-TBK1 (D1B4) Rabbit mAb (Cat# 3504S), anti-TRBP2 (D7C8K) Rabbit mAb (Cat# 62043S), anti-Dicer Ab (Cat# 3363S), anti-KHSRP Rabbit PolyAb (Cat# 55409-I-AP) were obtained from Cell Signaling (Danvers, MA, USA), anti-LGP2 Rabbit PolyAb (Cat# 11355-1-AP), anti-KHSRP Rabbit PolyAb (Cat# 55409-I-AP), anti-MCCC2/MCCB PolyAb (Cat# PA5-83410) were obtained from Thermo Fisher Scientific, Anti-PCCA antibody (Cat# AB154254) was obtained from Abcam and Anti-β-actin antibody (Cat# A5316) was purchased from Sigma Aldrich. Protein bands were detected using the ECL Prime Western Blotting Detection Reagent (Cytiva Life Sciences, Marlborough, MA, USA) with the Azure 300 (Azure Biosystems, Dublin, CA, USA). Band intensities were analyzed using Fiji (NIH ImageJ; http://rsbweb.nih.gov/ij/).

### RNA extraction and real-time reverse transcription PCR

Total cellular RNA was isolated from cells using the RNeasy isolation kit (Qiagen, Germantown, MD). The cDNA was synthesized from total RNA using TaqMan reverse transcription reagents (Thermo Fisher Scientific) with random hexamer primer, according to the manufacturer’s instructions. The expression levels of mRNA of gene of interest were measured using quantitative RT-PCR on a CFX96 real-time system (BioRad, Hercules, CA); the two-temperature cycle of 95 °C for 15 s and 60 °C for 1 min (repeated for 40 cycles) was used. Relative quantities of the transcript were calculated using the ΔΔCt method with GAPDH as a reference. Normalized samples were expressed relative to the average ΔCt value for controls to obtain relative fold change in expression levels. The probes specific to each gene were purchased from Applied Biosystems (Thermo Fisher Scientific). Probes for *IFNA1*(Hs04189288_g1), *IFNA2*(Hs00265051_s1), *IFNA4*(Hs01681284_sH), *IFNA5* (Hs04186137_sH), *IFNA6* (Hs00819627_s1), *IFNA7* (Hs01652729_s1), *IFNA8* (Hs00266883_s1), *IFNA13* (Hs04190680_gH), *IFNA16* (Hs03005057_sH), *IFNA17* (Hs00819693_sH), *IFNL1* (Hs01050642_gH), *IFNB1* (Hs01077958_s1) and *GAPDH* (Hs99999905_m1) were purchased from Thermo Fisher Scientific.

### Construction of PC and LGP2 expression vectors

PC gene was purchased from Origine (Rockville, MD, USA), and then restriction site HindIII and EcoRV sites were induced by PCR with 5’-AGAAGCTTGCCGCCATGCTGAAGTTCCG-3’ and 5’-GTGATATCCTCGATCTCCAGGATGAGGTC-3’. PCR product was cloned and inserted into pCMV-myc HA (TakaraBio, Cat# 631604) or pcDNA™3.1/V5-His A (Thermo Fisher, Cat# 43-0037) expression vectors using standard molecular biology techniques(32). Lysine at amino acid residue 912 of PC was mutated to Alanine (K912A) using sense and antisense oligo 5’-GTCACTCCCAGTTCCGCGATCGTGGGCGACCTG-3’ and 5’-CAGGTCGCCCACGATCGCGGAACTGGGAGTGAC-3’ respectively. The site direct mutagenesis was constructed using QuickChang II XL-Site-Directed Mutagenesis kit (Agilent, Santa Clara, CA, USA, Cat# 200521) (33). Lysine at amino acid residue 1144 of PC was mutated to Alanine (K1144A) using sense and antisense oligo 5’-TGTGCGTGTTGTCTGCAATGGCGATGGAAACTGTAGT -3’ and 5’-ACTACAGTTTCCATCGCCATTGCAGACAACACGCACA -3’, respectively.

The His-tagged or hemagglutinin (HA)-tagged LGP2 expression vectors were constructed as follows. LGP2-encoding complementary DNA (cDNA) was synthesized from the total cellular RNA of MDMs using the Superscript First-Strand System for RT-PCR (Thermo Fisher Scientific). The reverse-transcribed cDNA encoding LGP2 was subcloned into pCRXLII (Thermo Fisher Scientific). Then restriction site HindIII and EcoRV sites were induced by PCR with 5’-AGTTAAGCTTATGGAGCTTCGG -3’ and 5’-GCTGGATATCTGTCCAGGGAGAG -3’. PCR product was cloned and inserted into pCMV-myc HA (TakaraBio, Cat# 631604) or pcDNA™3.1/V5-His B (Thermo Fisher, Cat# 43-0037) expression vectors using standard molecular biology techniques (32). Histidine at amino acid residue 576 of LGP2 was mutated to Serine (H576S) using sense and antisense oligo 5’-GGTGGAGGGCACCAGCCATGTCAATGTG-3’ and 5’-CACATTGACATGGCTGGTGCCCTCCACC-3’, respectively. Phenylalanine at amino acid residue 601 and Tryptophan at amino acid residue 604 of LGP2 was mutated to Serine (F601S/W604S) using sense and antisense oligo 5’-CATCAACAAAGTCTCCAAGGACAGTAAGCCTGGGGGTGTC-3’ and 5’-GACACCCCCAGGCTTACTGTCCTTGGAGACTTTGTTGATG-3’, respectively.

Lysine at amino acid residue 634 of LGP2 was mutated to Glutamic acid (K634E) using sense and antisense oligo 5’-AAGCTGCCAGTGCTCGAAGTCCGCAGCATG-3’ and 5’-CATGCTGCGGACTTCGAGCACTGGCAGCTT -3’ respectively. The insertion was confirmed by Sanger DNA sequencing with BigDye version 3.0 (Applied Biosystems). Plasmid was screened by Sanger DNA Sequence and then intended plasmids were purified for transfection using EndoFree plasmid Maxi preparation kit (Qiagen, Germantown, MS, USA, Cat#12362).

### miRNA, siRNAs and Transfections

miRNA mimics and siRNA were synthesized by Thermo Fisher Scientific. miRNA mimic negative controls were obtained from Thermo Fisher Scientific (control 1: Cat # 4464058, control 2: Cat# AM17110) and GE-Healthcare (Chicago, IL) (control 3: Cat# CN0010000-01-05). An siRNA negative control was purchased from Thermo Fisher Scientific. siRNAs or miRNAs were transfected into cells using Lipofectamine RNAiMAX (Invitrogen, Thermo Fisher Scientific) according to the manufacturer’s protocols. DNA transfections were conducted with Lipofectamine 2000 (Invitrogen, Thermo Fisher Scientific) for MDMs or *Trans*IT-293 (Mirus Bio LLC,) for 293T or Hek cells, according to the manufacturers’ protocols. All synthesized DNA and RNA were obtained from IDT (Newark, NJ, USA).

### Isolation of recombinant proteins

The PC and LGP2 (Wt and mutant) proteins induced in 293T cells were isolated from cell lysate using Pierce ^TM^ IP cell lysis buffer (Thermo Fisher Scientific, Cat# 87788) with 5% glycerol, or with hypotonic buffer (50mM HEPES, 10mM KCl and 500mMEDTA) with protease and phosphatase inhibitors with EDTA. His-tagged PC and LGP2 proteins (Wt and mutant) were purified on a Ni-NTA column using His Pur ^TM^ Ni-NTA resin (Thermo Fisher scientific, Cat# 88222) via the carboxyl-terminal or N-terminal hexa histidine tag. HA-tagged recombinant proteins were affinity purified using HA-tagged specific monoclonal antibody conjugated on an agarose gel. After washing residual impurities, bound HA-tag proteins were eluted off the affinity column by 0.1M glycine (pH2) followed by neutralization with Tris (pH8.5).

### Cytosolic fraction

MDMs cytosolic fraction was prepared using A3G lysis buffer (50 mM HEPES (pH7.3)-125 mM mM NaCl-0.2% NP40). Briefly, 10 x 10^6^ MDMs were washed with cold PBS at 500 xg for 5 min and then cell pelleted were resuspended in A3G buffer containing protease and phosphatase inhibitor with EDTA (Thermo Fisher Scientific) at 10 x 10^6^/mL. Cell suspension was incubated on ice for 10 min and then lysed using a Dounce homogenizer with 30 strokes. After cell lysis was confirmed by microscope, entire cell lysate was centrifuged at 20,000 xg for 10 minutes, and the supernatants were collected as cytosolic fraction and stored at -80°C until use.

### Pull-down assay of miRTC14

MDMs were used to get cell lysate using hypotonic buffer in nuclear extraction kit (Cat# 40010, Active Motif, Carlsbad, CA) and pelleted cell debris by centrifuging at 14 000g for 10 min at 4°C. A total of 1 mg of cytoplasmic proteins were incubated with 100 µl of Dynabeads M-280 streptavidin (Invitrogen, Thermo Fisher Scientific) conjugated with 2 nmol 5′ biotinylated miRTC14 and 2 nmol 3′ biotinylated miRTC14.

Cell-lysates were pre-cleaned to remove non-specific binding to the beads. Pre-cleaned cell lysates were processed under two experimental conditions, such as no competitor condition and competitor condition. For non-competitor condition pre-cleaned cell lysate was incubated with 100 µl of miRTC14 conjugated dynabeads. For competitor condition 20nm non-biotinylated miRTC14 was incubated with condition pre-cleaned cell lysate prior to mixing with100 µl of miRTC14 conjugated dynabeads. Then both non competitor and competitor samples were incubated for 2-4 h at 4 °C. Precipitated oligonucletide–protein complexes were washed extensively in protein binding buffer (20 mM HEPES, 50 mM KCL, 10% glycerol, 5 mM MgCL2 and 1 mM dithiothreitol (DTT, Sigma-Aldrich). Proteins were eluted from the beads by boiling in NuPAGE lithium dodecyl sulfate (LDS) sample loading buffer (Thermo Fisher Scientific) and were separated on NuPAGE 4–12% Bis-Tris gel Thermo Fisher Scientific) and stained using Silver Quest (Invitrogen). Individual bands were excised from the gel, digested using trypsin and analyzed by liquid chromatography and mass spectrometry (MS; LTQ XP, ThermoElectron, San Jose, CA, USA (34).

### Biolayer interferometry

Real-time binding assays of recombinant histidine tagged pyruvate carboxylase (PC) and LGP2 to miRNAs were characterized by biolayer interferometry (BLI) using an Octet R8 (Sartorius, NY, USA) with Octet Ni-NTA Biosensors (Cat # 18-5101) as previously described (35,36). All proteins and miRNAs were in PBS + 0.02% Tween-20 + 0.01% BSA including 1 mM DTT and 1 mM MgCl_2_ for protein stability. First, Octet Ni-NTA Biosensors were hydrated by immersing them Octet kinetic buffer for at least 10 min. The BLI protocol was as follows: 60 s baseline1, 900 s loading with 1 μM protein, 300 s baseline2, 600 s association with 10nM of miRCtrl, miRSX2, miRTC10 and miRTC14 and 600 s dissociation. The assays were performed on black bottom 96-well microplates (Greiner Bio-One 655209) in a total volume of 200 μL, at 30 °C with orbital shaking at 1000 rpm. Experiments were conducted with the software Octet Data Acquisition software (Sartorius, USA Kinetic binding parameters were calculated using Octet R8 Data Analysis 8.2 (Sartorius). Control experiments were conducted in buffer without proteins. Control data from reference sensors were used to correct for biosensor drift and other artifacts. After subtraction of reference biosensors, the binding curves were aligned to the X- and Y-axis and the association-dissociation inter-step curve in order to get a common baseline for the association and dissociation phases. BLI results were analyzed in Octet Analysis Studio 13 (Sartorius). If R^2^>0.95 and RSS is <3 considered as 1:1 fit. Binding kinetic parameters were obtained by globally fitting the data using a 1:1 binding model available in the analysis software.

### Immunofluorescence staining and confocal microscopy

MDMs were seeded onto 12 mm coverslip-inserted 12-well cell culture plates and transfected with 50 nM single-stranded miRTC14 as indicated. At 18 h post-transfection, cells were washed with PBS, fixed with 4% formaldehyde (Thermo Fisher Scientific) for 15 min at room temperature, and permeabilized with 0.1% Triton X-100 (Thermo Fisher Scientific) for 10 min. After blocking with blocking buffer for 1 h at RT, cells were then incubated with primary antibody against PC at 4°C overnight, followed by incubation with appropriate AF555-conjugated secondary antibodies for 1 h at room temperature. Mitochondria were stained using MitoTracker (Thermos fisher) according to the manufacturer’s instructions. Coverslips were mounted using antifade mounting medium with DAPI (Thermo Fisher). Imaging was carried out on a Zeiss Axio Observer.Z1 motorized LSM800 confocal microscope using a Plan-Apochromat 63x/1.40 oil objective (Zeiss, Oberkochen, Germany).

Colocalization analyses of the images were carried out using Fiji/Image J (2.16.0/1.54p) using the colocalization threshold plugin. Regions of interest (ROIs) were selected as green channel (MiRTC14 signalling), and colocalization between AF488-labeled mirTC14 and PC was quantified using Rcoloc coefficients. All imaging settings were kept consistent across samples for comparative analysis. The images shown are representative of three independent experiments.

### Enzyme-linked immunosorbent assay (ELISA)

The level of IFN-α, IFN-λ1 and IFN-β protein in culture supernatants was measured using Human IFN-alpha All Subtype Quantikine ELISA Kit (R&D Systems), Duoset human IFN-λ1 ELISA kit or human IFN-β ELISA kit (R&D Systems) according to the manufacturer’s protocol.

### Computational method for building the binding model between miRNA and PC/LGP2

Three miRNAs analyzed in this study—miRTC14, miRTC10, and miRSX2—are listed in Table 1. The double-stranded miRNAs mimic (ds-miRNAs by adding double U at each 3” end forming the overhang structure) were used in the miRNA-protein interaction prediction analysis. The potential miRNA binding protein used in this study was PC (UniProt ID: P11498). Docking predictions between double-stranded miRNAs and PC were performed using AlphaFold 3 (37) via the AlphaFold Server (https://alphafoldserver.com/) with default parameters (37), where five predicted models were returned for each miRNA-PC docking analysis.

**Table 1.**
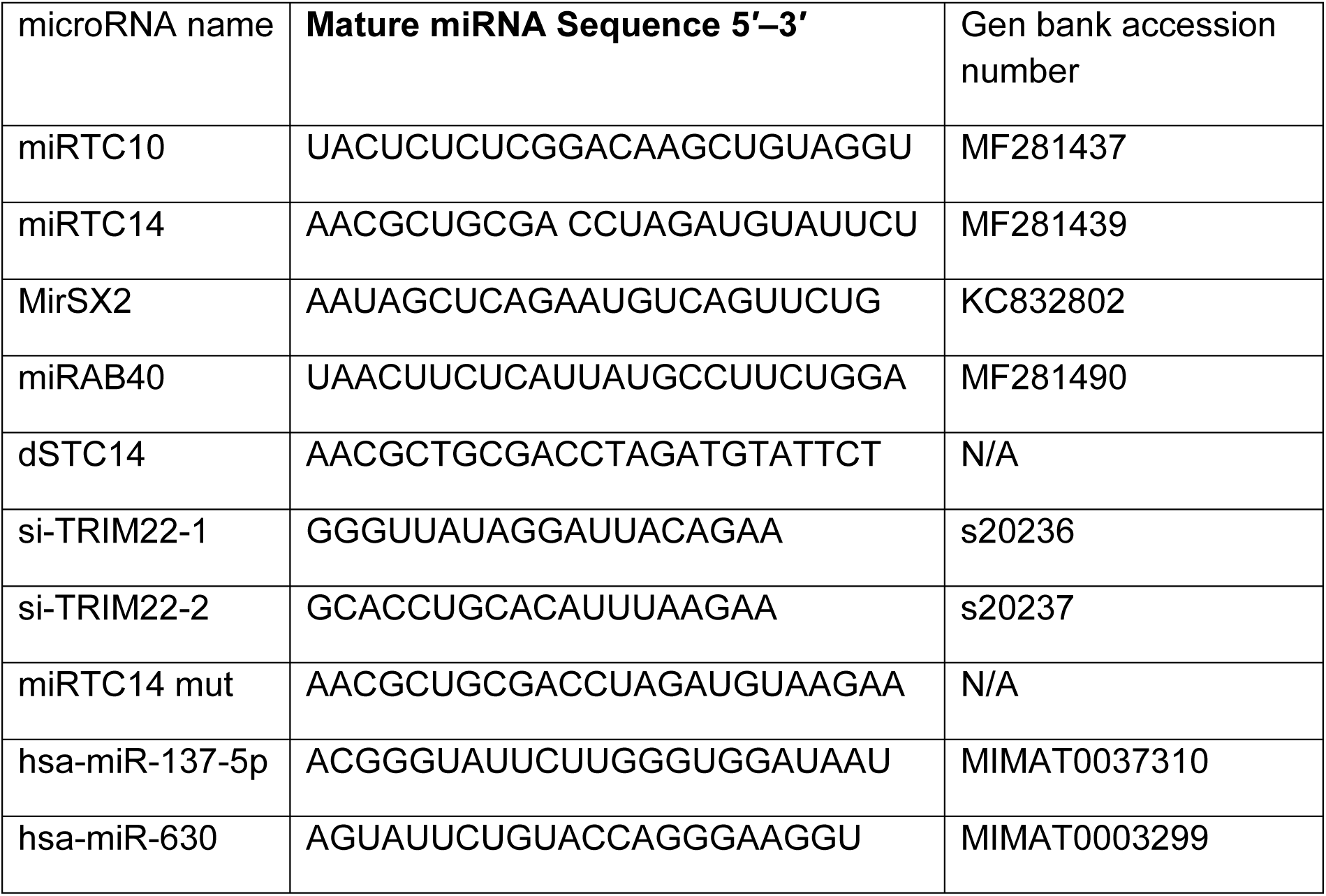
Sequences of miRNAs.

To identify the most favorable miRNA–PC interactions, the predicted docking models from alphaFold 3 were analyzed, and the dominant binding poses were selected. To compare the binding of miRNAs to PC, the interaction energy of each miRNA-PC complex predicted by AlphaFold 3 was calculated using FoldX (https://foldxsuite.crg.eu/) (38). The representative models were selected according to their FoldX-predicted binding energies, with the lowest binding energy values chosen as representative. To validate the predicted binding sites, *in-silico* mutagenesis was conducted by introducing mutations either on the key miRNA nucleotides or on the amino acid residues of PC. The effects of protein mutations on miRNA–PC interactions were evaluated using mCSM-NA (39) and PremPRI (40).

For building the miRTC14 and LGP2 (UniProt ID: Q96C10) binding models, two LGP2 models were constructed, depending on the capping loop conformations (41,42). For the closed capping loop LGP2 model, the binding complex with miRTC14 was predicted using AlphaFold 3. The prediction procedure followed the same workflow as that used for the miRNAs-PC complex. The LGP2 model with an open capping loop was generated by homology modelling using SWISS-MODEL (43), with the crystal structure of LGP2 (PDB ID: 9LOV) as the template. The three-dimensional (3D) structure of miRTC14 was predicted using AlphaFold 3. Protein-RNA docking was then performed with GRAMM (44,45) using the 3D structure of miRTC14 and the 3D model of LGP2 with an open capping loop as input structures.

The representative miRNA-PC and miRNA-LGP2 complex models were further refined using YASARA (46) to obtain more favorable conformations. 3D visualization of miRNA-PC interaction was generated using the open-source version of PyMOL (https://github.com/schrodinger/pymol-open-source), while the corresponding two-dimensional (2D) interaction diagram was generated using PDBsum1 (https://github.com/RomanLas/PDBsum1). Finally, experimental validation was performed to confirm computational predictions.

### Statistical analysis

Results were representative of at least three independent experiments. The values were expressed as mean and standard deviation (SD) of individual samples.

Intergroup comparisons were performed by one-way ANOVA with multiple comparison analysis or Student’s t-test using GraphPad Prism 9 (GraphPad, San Diego, CA, USA). p-Values less than 0.05 were considered statistically significant: * p < 0.05, ** p < 0.01, *** p < 0.001, ****p <0.0001, and p > 0.05 was considered not significant (ns).

## RESULTS

### Characterization of anti-HIV effect

In our previous studies, we showed that IL-27–differentiated T cells and macrophages induce a set of novel antiviral microRNAs (miRNA), including miRTC10, miRTC14, miRSX2, and miRAB40, which exhibit anti-HIV activity (25,26,47) and promote IFN production, although the underlying mechanism remained unclear. In this current study, we investigated the mechanism underlying the miRNA-induced IFNs. We first compared among the anti-HIV miRNAs. MDMs-transfected with individual miRNA mimic were infected with HIVAD8 (replication competent) or HIVLuc-VSV-G (pseudo-typed replication incompetent). HIV replication was then compared to that in control miRNA-transfected cells. This screening confirmed miRSX2, miRAB40, miRTC10, and miRTC14 as antiviral miRNAs (Supplementary Fig. S1A and 1B). Dose–response analysis of anti-HIV activity using HIVLuc-VSV-G revealed that all four miRNAs suppressed HIV replication in a dose-dependent manner (Fig.1A-D).

**Figure 1:**
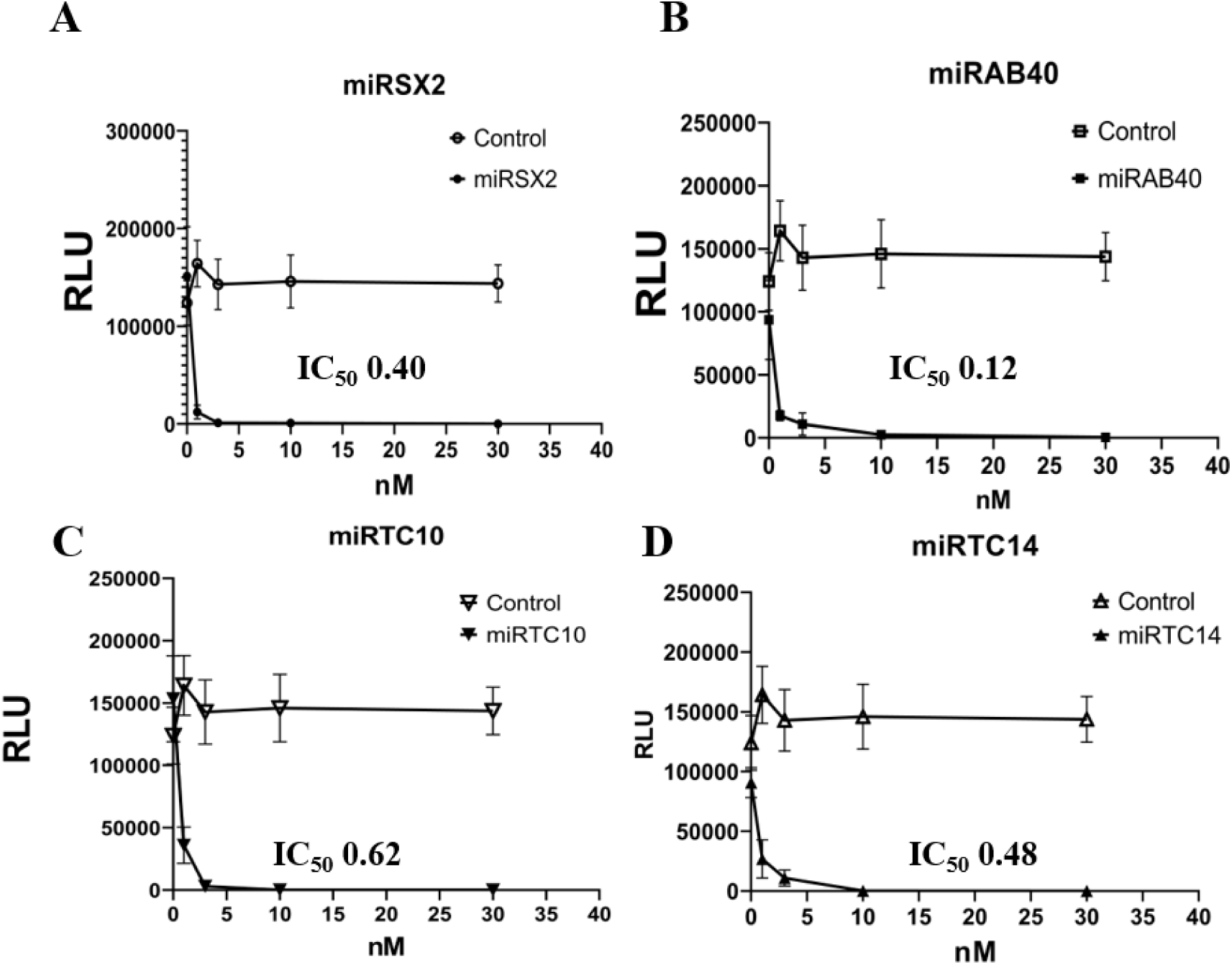
miRNA dose-dependently suppressed HIV infection. Different dose of four miRNA mimics (miRSX2 (**A**), miRAB40 (**B**), miRTC10(**C**), and miRTC14(**D**),) were transfected into MDMs, and then cells were infected with HIVLuc-VSVG. Anti-HIV activity was assessed by Luciferase activity as described in Materials and Methods. The 50% inhibitory dose (IC₅₀) of each miRNA was then determined and shown within the figure. Results show means ± SD from one representative data from four independent assay.

### Characterization of IFN inducing activity

To assess whether the anti-HIV effect in these four miRNAs was mediated via IFNs, an IFN absorption assay was conducted using B18R, a soluble vaccinia virus-derived IFN type-I receptor blocker that suppresses the effect of IFN-α/β (48). As a positive control for the B18R effect, HIV-infected cells were cultured with IFN-α. In the presence of B18R, IFN-mediated HIV inhibition was suppressed to 100%, and HIV replication in miRNA-treated cells was also restored. However, the level of replication recovery differed among the miRNAs (Supplementary Fig. S2). Inhibited HIV replication in miRTC14-transfected cells recovered to 79 ± 3.6%, whereas replication in miRSX2- and miRTC10-treated cells recovered to 25 ± 6.2% and 39 ± 3.8%, respectively, compared to control miRNA (miRCtrl)-transfected cells. These results suggest that type I IFN induction is involved in the anti-HIV effect of each miRNA; however, the dependency on IFN for inhibition may differ for each miRNA.

To elucidate whether IFNs were produced in culture supernatants after miRNA transfection, IFN-α, -β and -λ1 in the culture supernatants were quantified using specific ELISA kits. Since human IFN-α contains 12 subtype proteins in 13 functional genes (49,50), an all-subtype IFN-α ELISA kit was used in this study (Fig. 2A and Supplementary Table S1). We also investigated IFN-λ1 induction because this response is considered as an innate immune response to small-RNA transfection (24). The order of IFN production from high to low sequence was as follows: miRTC14 > miRSX2 > miRTC10 > miRAB40. Although amounts of IFN-α/β production from miRTC14, miRSX2 and miRTC10 were more than 10-fold higher than that observed from miRCtrl transfected cells. Of note IFN-λ1 secretion was selectively elevated (∼10-fold) by miRTC14 compared to miRCtrl-transfected cells, but not by miRTC10, or miRAB40 (Fig. 2A-C and Supplementary Table S1). Thus, miRTC14 was the most IFN-inducing miRNA among the four.

**Figure 2:**
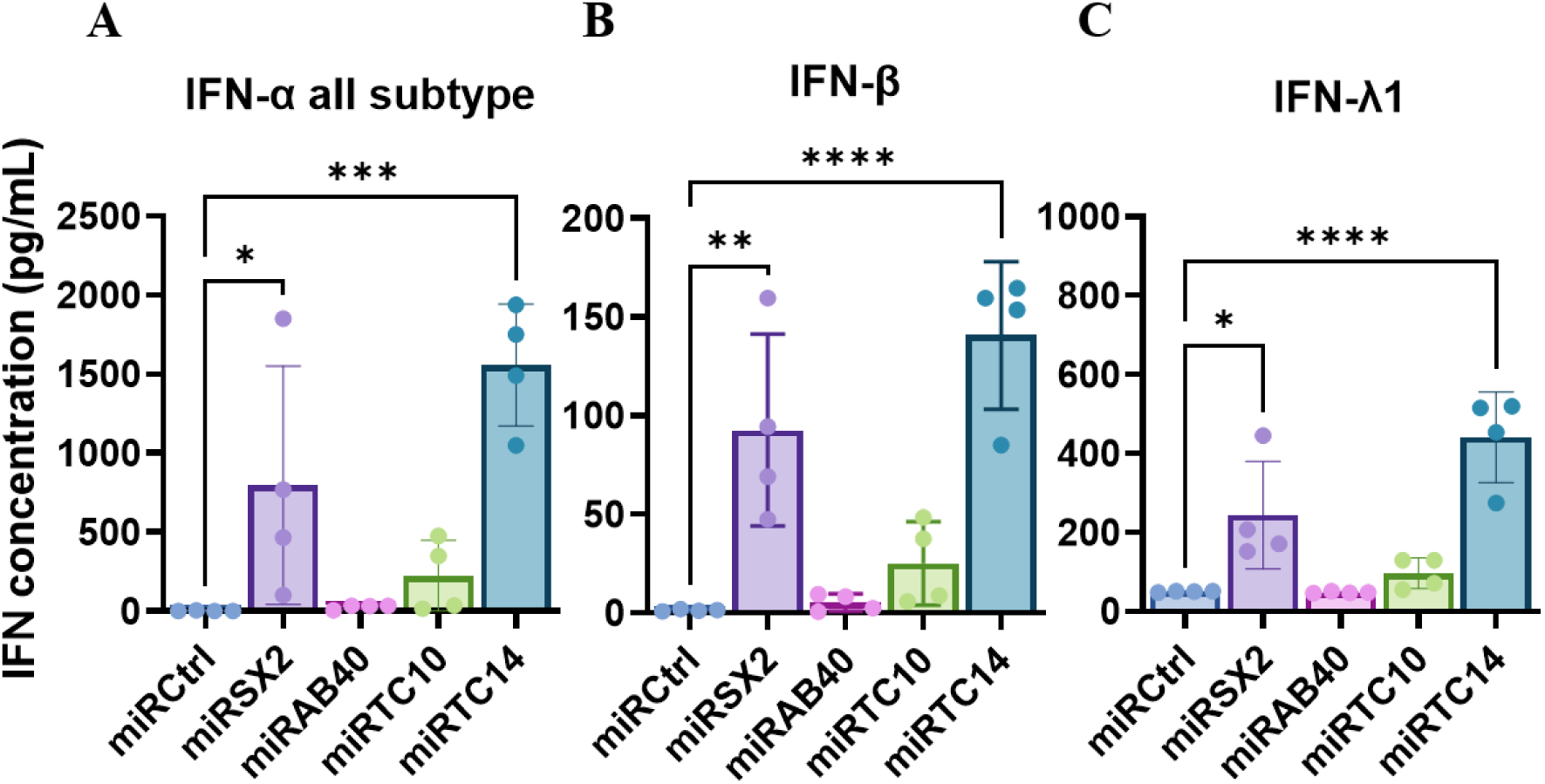
Protein levels of Type I and III interferon after anti-HIV microRNA transfection. MDMs were transfected with 10 nM of miRCtrl, miRSX2, miRAB40, miRTC10, or miRTC14 for 72 hours, and then, cell-free supernatants were collected. Concentrations of IFN-α, β, and λs were quantified using each IFN ELISA kit. IFN-α was detected using the all-subtype IFN-α kit. The detection limits of IFN-α, β, and λ1 were 1.25, 1.2, and 15.6 pg/mL, respectively. The bar graphs show data the results from 4 donors for IFN-αs **(A)**, IFN-β **(B)** and IFN-λ1 **(C).** *, **, and *** indicate *p* values are < 0.05, < 0.01, and < 0.001, respectively.

To determine which IFN-α gene subtype was activated by each miRNA transfection, qRT-PCR was performed using total cellular RNA with a total of 11 IFN-α subtype probes. One of these probes, *IFNA4*, is highly similar to *IFN10A*, with 98.4% identity. Therefore, we considered the result from *IFNA4* expression to include *IFNA10* expression. miRTC14 consistently induced *IFNA1*, *IFNA2*, *IFNA4/A10*, *IFNA5*, *IFNA7*, *IFNA8*, *IFNA13*, and *IFNA14* regardless of donor (Fig. 3A-J). Similar to *IFNA* data, miRTC14 also induced *IFNB1* (Fig. 3K) and *IFNL1* (Fig. 3L). The second most potent IFN inducer, miRSX2, prompted the induction of the same subtypes, except for *IFNA4/A10*, however, gene activation was dependent on the donor. miRTC10 transfection activated a lower level of multiple IFN genes, and donor dependency was observed. These results indicate that IFN genes may be activated differentially by miRNA, suggesting that IFN induction may be RNA sequence dependent.

**Figure 3:**
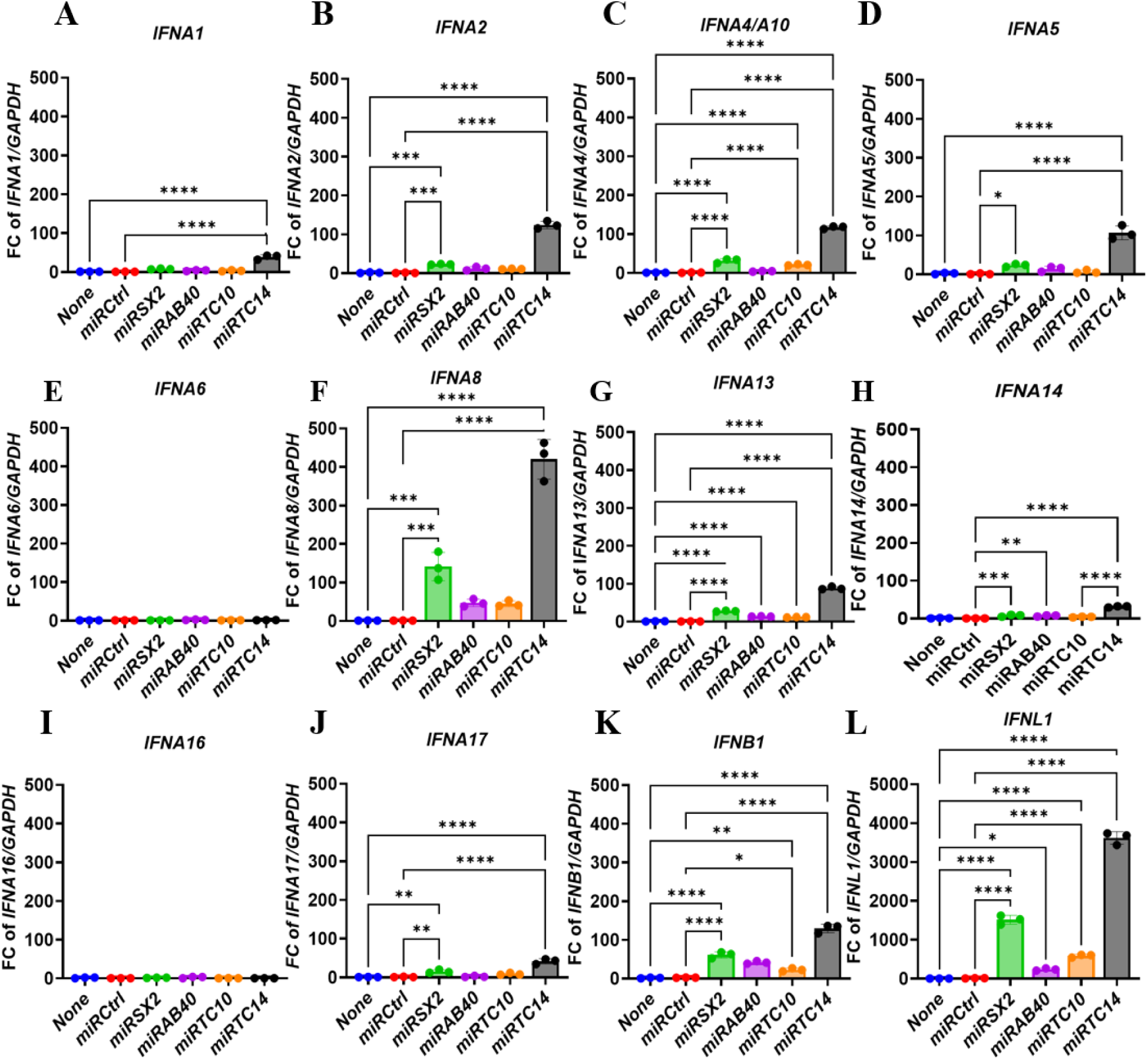
Characterization of IFN gene activation by microRNAs. Total cellular RNAs from MDMs from three independent donors were extracted after transfection with or without miRNA mimics. Then real-time qRT-PCR was performed using specific probes for each miRNA. GAPDH was used as an internal control, and each IFNs expression was calculated delta-Ct (ΔCt) using Ct values in each miRNA mimic and then gene expression is presented as relative expression units compared with un-transfected cells (none) after normalization to GAPDH. Results show the fold change of mean ± SD (n = 3) of *IFNA1*(**A**), I*FNA2*(**B**), *IFNA4/A10* (**C**), *IFNA5* (**D**), *IFNA6* (**E**), *IFNA8* (**F**), *IFNA13*(**G**), *IFNA14*(**H**), *IFNA16*(**I**), *IFNA17*(**J**), *IFNB1*(**K**), *IFNL1*(**L**). *, **, and *** indicate p values are < 0.05, < 0.01, and < 0.001, respectively.

Since miRTC14 was identified as the most potent IFN inducer, we next examined the contribution of its individual strands—passenger (p.miRTC14) and guide (g.miRTC14)—for IFN induction. Each strand was independently transfected into MDMs, and IFN expression was quantified by qRT–PCR. Double-stranded miRTC14 served as a positive control. Our results demonstrate that both double-stranded miRTC14 and its individual passenger and guide strands can induce the expression of *IFNA2*, *IFNA8*, *IFNA13*, and *IFNL1* (Supplementary Fig. S3A-D). However, the magnitude of IFN induction by the single-stranded forms (p.miRTC14 and g.miRTC14) was significantly lower than that observed with double-stranded miRTC14. Although both single strands retain ability to induce IFN, their reduced induction highlights the importance of duplex integrity. The enhanced activity of the double-stranded form likely reflects increased structural stability compared to single-stranded microRNAs and improved recognition by RNA sensors, facilitating more efficient engagement of downstream signalling partners. This is in line with established principles of innate immunity, in which double-stranded RNA serves as a key molecular pattern associated with viral infection (51,52).

### Identification of the mechanism of IFN gene activation

As shown in Table 1, the four miRNAs exhibited a lack of sequence similarity but induced IFNs even though the induction level was diverse. To define whether a given common target gene(s) is involved in the induction of IFNs, potential target for each miRNA was analysed using four prediction tools (miRDB, MR-microT, miRanda, and Target Scan) (53). A Venn diagram analysis was conducted to define whether common target gene(s) present (Supplementary Fig. S4 and Supplementary Table S2). Common target genes for four miRNAs were selected for further analysis, using the Database for Annotation, Visualization, and Integrated Discovery DAVID (54) and Metscape (55). Functional annotation analysis for these 11 genes, were not classified in IFN-induction functional category (Supplementary Fig. S5). Therefore, we concluded that the miRNA-mediated IFN induction was unlikely to result from the canonical gene-silencing effect. Instead, the IFN responses were more likely attributable to off-target effects by miRNAs transfection. Since some, but not all miRNAs induced the IFNs induction, it was considered that these responses might be a nucleotide sequence dependent, which is different from an innate immune response mediated via RIG-I like receptors, which is sequence independent immune reaction. These findings raise the possibility that sequence-selective intracellular RNA recognition mechanisms contribute to IFN induction.

### Pyruvate carboxylase and LGP2 are binding partners of miRTC14

To determine whether a specific protein binds to miRNAs, employing miRTC14, the most IFN-inducing miRNA, in a competitive pulldown assay (56). As illustrated in a scheme in Supplementary Fig. S6, MDMs whole-cell lysate was split into two fractions, and one fraction (control reaction) was mixed with biotinylated miRTC14 (bio-miRTC14), and another fraction (competitive pull down) was mixed with bio-miRTC14 and 10-fold molar excess of unlabelled miRTC14. The protein mixture was incubated and then pull-down using streptavidin-conjugated beads. After washing, bound proteins were separated on SDS-PAGE and visualized using the SimplyBlue Safe staining. Band 1, band 2, band 3 and band 4 in the control lane were consistently diminished in the competition lane, but other bands were not significantly changed (Fig. 4A, Supplementary Fig. S7A-B). These results suggested that the vanished bands in the gel were potential miRTC14-binding proteins, thus the bands were excised and then subjected for protein mass analysis.

**Figure 4.**
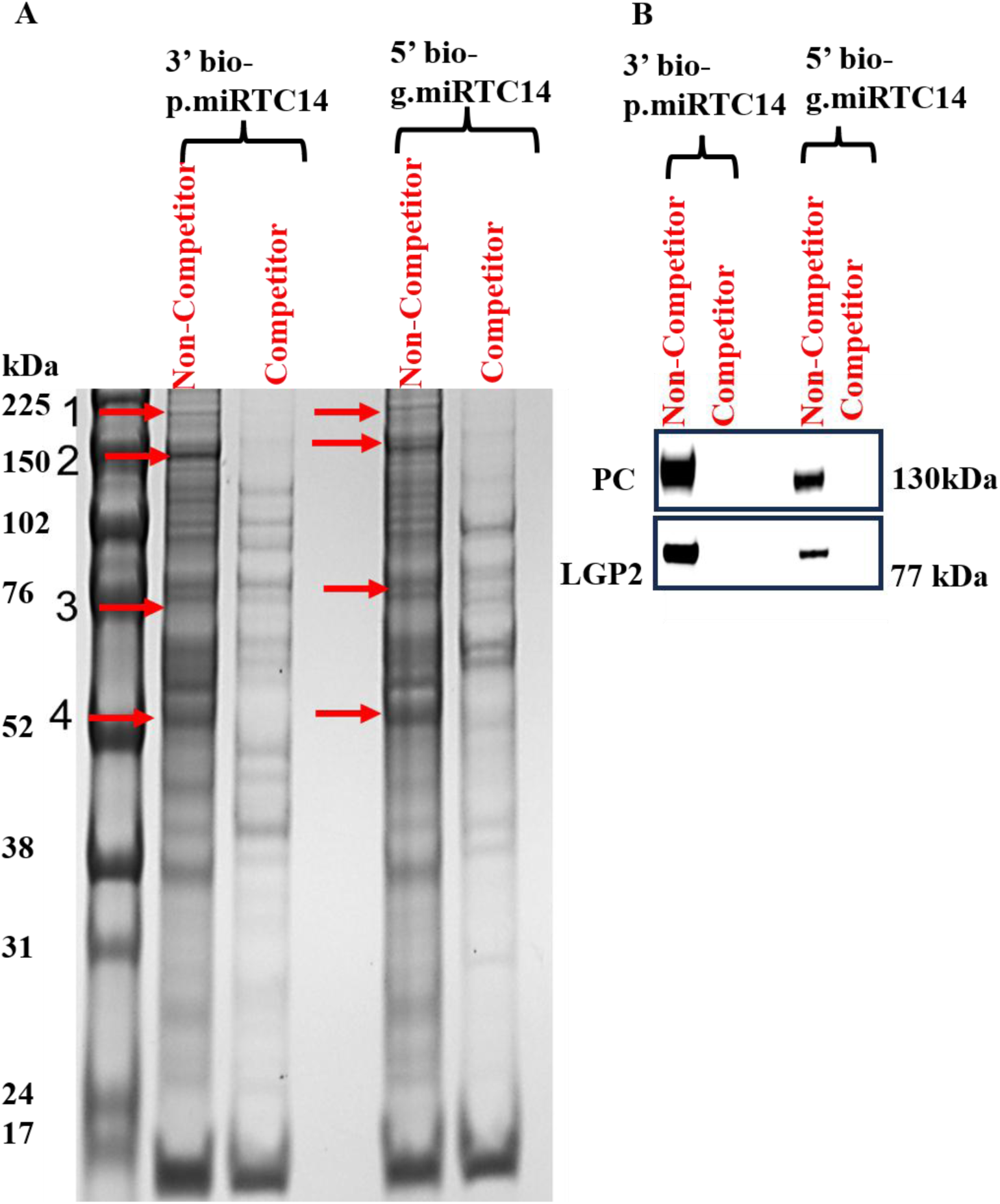
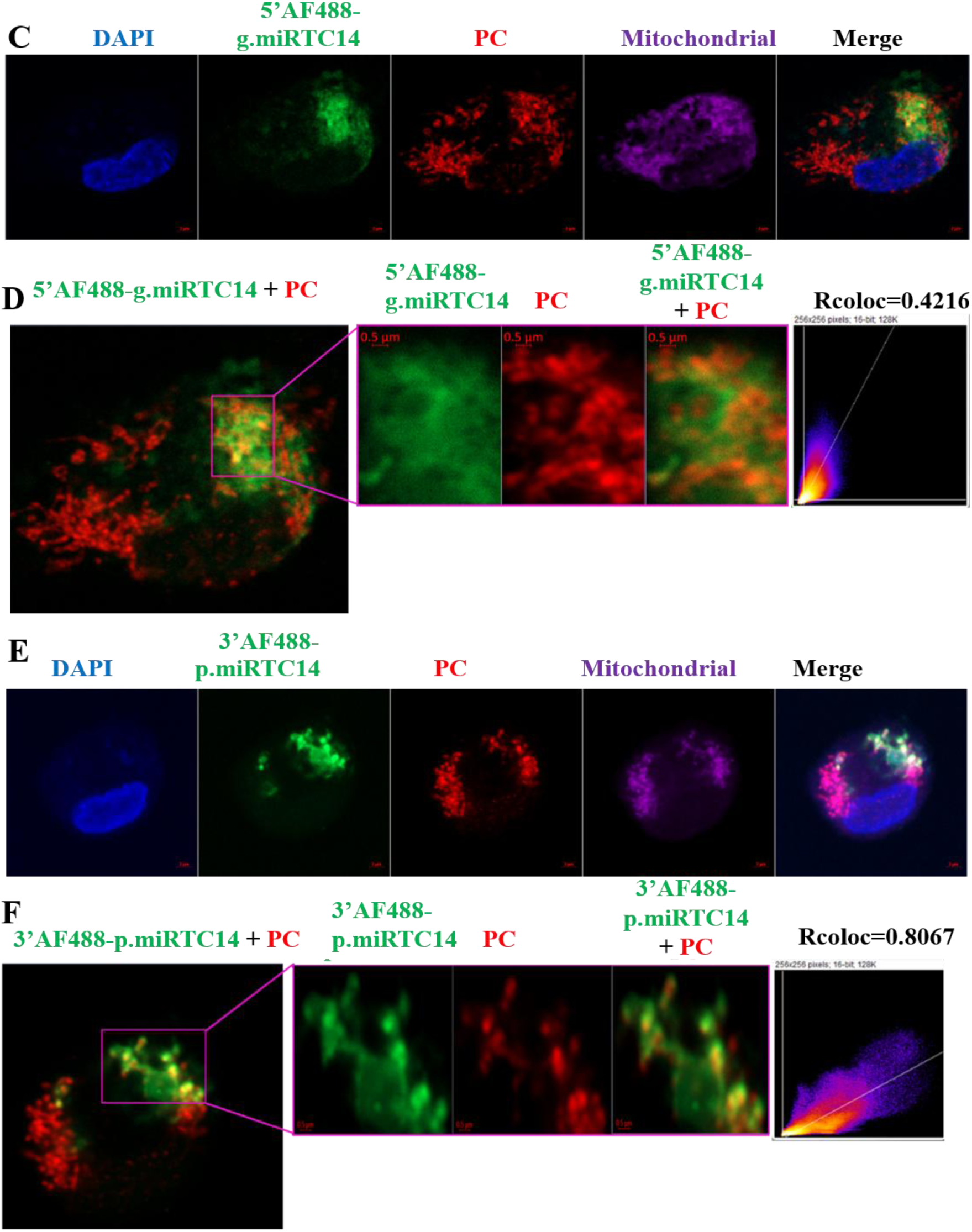
Identification of miRTC14-binding proteins. **(A)** SDS-PAGE of proteins recovered from pull-downs beads were visualized by Coomassie staining. Bands 1–4 present in the control lane were markedly reduced in the competitive condition, indicating specific miRTC14-binding proteins. These bands were exercised and subjected to mass spectrometry for identification. (**B**) Western blot analysis of proteins to confirm miRTC14 bound proteins using anti-PC antibody and anti-LGP2 antibody. (**C & E**) Confocal microscopy was used to determine the subcellular localization of the transfected guide strand (**C**) or passenger strand (**E**) of miRTC14 (green) and PC (red). MDMs were seeded on 12 mm coverslip-inserted 12-well plates and transfected with 50 nM guide or passenger-stranded miRTC14. At 18 h post-transfection, cells were fixed and stained with anti-PC antibody and MitoTracker (purple); nuclei were counterstained with DAPI (blue). (**D & F**) Colocalization analysis was performed using Fiji within the indicated rectangular regions of interest. Rcoloc coefficients are reported to quantify the degree of colocalization between the 5′ AF488–labeled miRTC14 guide strand (**D**) or 3’ AF488-labeled passenger strand (**F**) of miRTC14 (green) and PC (red).

Results of the mass spectrometry analysis showed that these bound proteins included known RNA binding protein such as Transactivation Response Element RNA-binding Protein (TRBP2) (57), 3-methylcrotonyl-CoA carboxylase B (MCCB) (58), Protective protein/Cathepsin A (PPCA) (59), Dicer (60), KH-type splicing regulatory protein (KHSRP) (61), and DHX58/LGP2 (16) (Supplementary Fig. S7C). Surprisingly, we found a metabolic enzyme, pyruvate carboxylase (PC) (16,17). To confirm protein identification, WB assay was conducted. As demonstrated in Fig. 4B and Supplementary Fig. S7C, the presence of each candidate confirms protein in the corresponding gel band. Dicer, TRBP2 are known miRNA binding proteins (17,62,63), LGP2 is known double-stranded cytosolic RNA sensor, which regulates RNA-mediated innate immune response (64,65). Given that PC is a mitochondrial protein, it prompted us to investigate whether the transfected miRTC14 forms a complex with PC in the mitochondria.

To elucidate the distribution, a confocal microscopic analysis was conducted using MDMs. In this study, the synthesis of fluorophore-conjugated duplex miRTC14 was not successful. Consequently, we opted to utilize AF488-tagged single-stranded miRTC14 as an alternative. Confocal imaging exhibited that both the 5′ and 3′ AF488–labelled guild and passenger strands of miRTC14 localized nearby mitochondria-associated PC in MDMs (Fig. 4C and E). The green guide strand signal overlapped with red PC staining, especially within mitochondrial areas marked by MitoTracker. Quantitative analysis using Fiji confirmed spatial overlap, with Rcoloc coefficients of 0.4216 for the guide strand (Fig. 4D) and 0.8067 for the passenger strand (Fig. 4F). As expected, PC predominantly expressed in mitochondria (Fig.4C and E). These findings indicate that miRTC14 polarized near mitochondria and may directly or indirectly interact with PC in MDMs.

### Measurement of microRNAs binding to the recombinant PC and LGP2 by biolayer interferometry assays

To determine whether the interaction between PC/LGP2 and miRTC14 was direct or indirect, we performed binding analysis using recombinant proteins with bio-layer interferometry technology (BLI) on an Octet R8 instrument. To avoid potential interference resulting from improper orientation or steric hindrance associated with immobilization through amino groups, purified recombinant 6x His-tagged PC and LGP2 proteins (Supplementary Fig. S8A-B) were immobilized on Ni-NTA sensor chips, allowing miRNAs to flow freely and interact with the proteins. We tested miRCtrl, miRSX2, miRTC10, and miRTC14 against immobilized PC or LGP2. As shown in Fig. 5A and C and Supplementary Table S3, the dissociation constants (K_off_) for miRTC14 binding to PC (1.09E-05 ±1.97E-08) were >100-fold tighter than those observed for miRTC10 (9.44E-03 ± 5.70E-08), indicating miRTC14 exhibited stronger binding than miRTC10. Furthermore, specific binding of miRTC14 to PC (K_off_ = 1.09E-05 ±1.97E-08) were higher than that observed with LGP2 and miRTC14 (K_off_ = 1.91E-04 ± 3.16E-08) (Fig. 5A-B), indicating that sequence variations significantly influence the binding. In contrast LGP2 demonstrated stronger binding to miRTC10 (K_off_ = 1.52E-05 ± 6.18E-07), compared to miRTC14 (K_off_ = 1.91E-04 ± 3.16E-08) (Fig. 5B and D). miRCtrl and miRSX2 showed no detectable binding to either PC or LGP2 (Fig. 5E). Of note, although miRSX2 is an IFN-inducing miRNA, it did not bind to PC.

**Figure 5.**
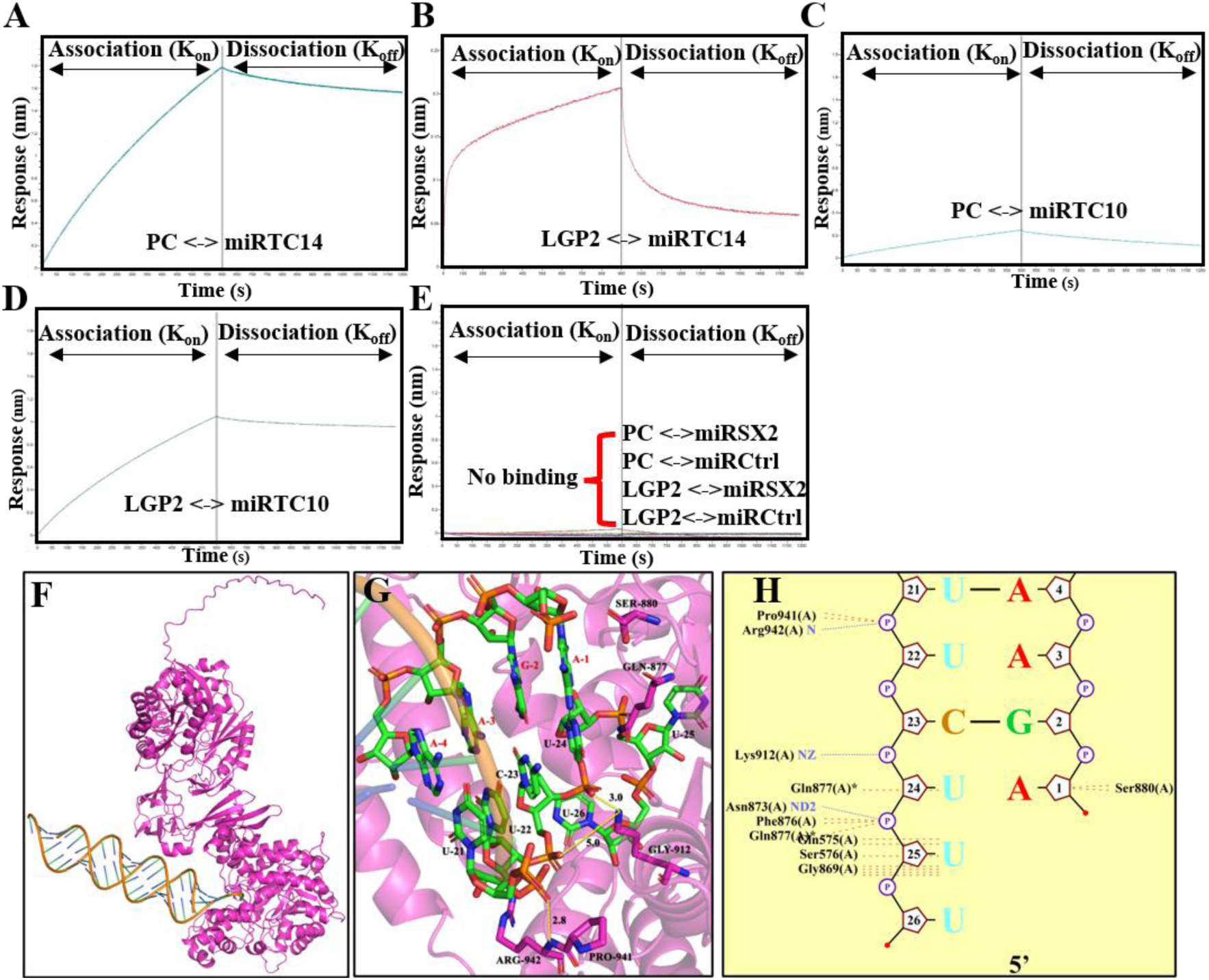
Direct binding analysis of miRNAs to PC and LGP2. To determine whether the interaction between miRTC14 and the RNA-binding proteins PC and LGP2 is direct, binding kinetics were assessed using BLI on an Octet R8 instrument. Purified recombinant 6×His-tagged PC and LGP2 proteins were immobilized on Ni–NTA sensor chips to ensure proper orientation and minimize steric hindrance, while miRNAs were allowed to freely associate in solution. miRCtrl, miRSX2, miRTC10, and miRTC14 were evaluated for binding to immobilized PC or LGP2. Control experiments were conducted in buffer without proteins. PC showed strong binding to miRTC14 (**A**) and less binding to miRTC10 (**B**). LGP2 bound to miRTC14(**C**) and miRTC14 (**D**). miRSX2 and miRCtrl showed no binding to both PC and LGP2 (**E**). Binding kinetic parameters were obtained by globally fitting the data using a 1:1 binding model available in the analysis software to obtain kinetic parameters k_on_ and k_off_. The experimental curves are shown as solid lines. AlphaFold 3 was used to predict docking models for each miRTC14–PC complex (**F, G**). Predicted in main interaction nucleotides in each strand of miRTC14 and PC (**H**).

To further evaluate the interaction, we performed docking model analyses as described in the Materials and Methods. AlphaFold 3 was used to predict docking models for each miRTC14–PC and miRTC14–LGP2 complex. Among the top predicted models, miRTC14 consistently exhibited similar binding poses within the putative PC binding site (Fig. 5F-H). These findings suggest that miRTC14 adopts more stable and consistent docking conformations at the predicted binding site. The interaction energy was further estimated using FoldX, and the optimal predicted values for each miRNA–PC complex were selected for comparison (Supplementary Table S4). Notably, the miRTC14–PC complex exhibited the most favourable binding energy of -8.15 kcal/mol evaluated by FoldX, indicating a stronger binding. Similarly, miRTC14 showed favorable binding to LGP2, however the binding sequences are different from PC-miRTC14 interactions (Supplementary Fig. S9, A to F).

### PC and LGP2 play a key role in miRTC14 mediated IFN induction

LGP2 plays a key role in RIG-I–like receptor–mediated antiviral responses (8,12). Given that miRTC14 shows stronger and tighter binding to PC, we focused on the role of PC and LPG2 in the miRTC14—— mediated IFN induction. To evaluate the contribution of each protein, each protein was individually knocked down in MDMs using specific siRNAs. Cells were transfected with si-control (si-Ctrl), si-PC, or si-LGP2, followed by transfection with miRCtrl or miRTC14. IFN induction was then quantified by qRT-PCR. Consistent with previous results (Fig. 3), miRTC14 robustly induced the expression of *IFNA2*, *IFNA8*, *IFNA13*, and *IFNL1*. However, this induction was partially reduced in both si-PC–treated (Fig. 6A-D) and si-LGP2– treated cells (Fig. 6E-H). These findings indicate that both PC and LGP2 are required for miRTC14-mediated IFN induction. Nevertheless, it remains unclear whether this effect results from the independent contributions of each protein or from a coordinated and cooperative mechanism.

**Figure 6.**
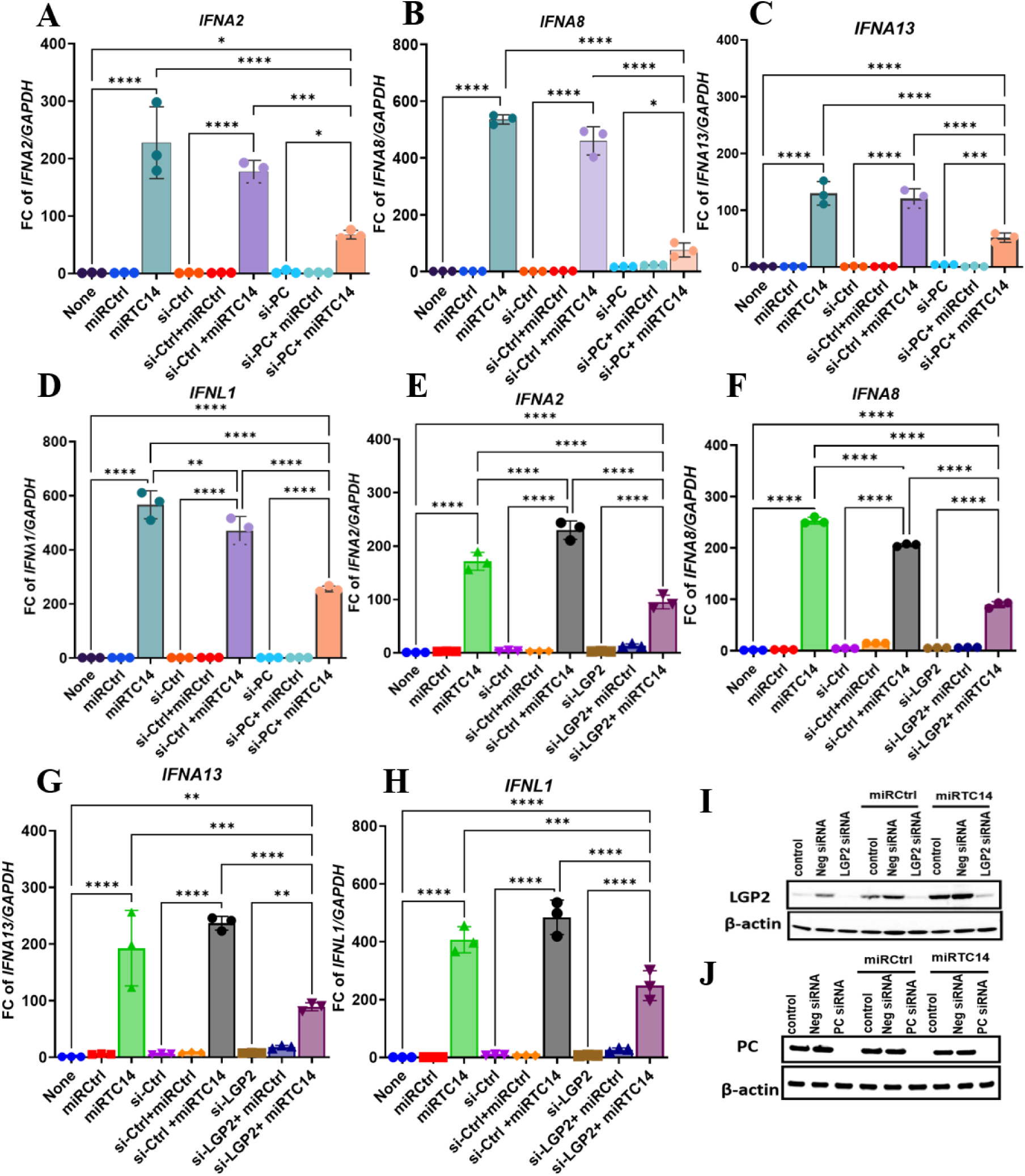
Silencing PC and LGP2 disrupted miRTC14-mediated IFN induction. To assess the contribution of PC and LGP2 to miRTC14-induced IFN responses, MDMs were subjected to siRNA-mediated knockdown using control siRNA(si-Ctrl), PC siRNA (si-PC), or LGP2 siRNA (si-LGP2). Cells were subsequently transfected with control microRNA (miRCtrl) or miRTC14, and IFN expression was quantified by qRT–PCR. Gene expression is presented as relative expression units compared with no-transfection after normalization to GAPDH. The IFN induction by miRTC14 was partially attenuated in both si-PC–treated cells **(A–D)** and si-LGP2–treated cells **(E– H).** Results show mean ± SD (*n* = 3). *, **, and *** indicate *p* values are < 0.05, < 0.01, and < 0.001, respectively. (**I-J**). Silencing LGP2 and PC protein expressions. MDMs were transfected with si-LGP2 (**I**) or si-PC **(J**) and knockdown efficiency of LGP2 following siRNA treatment was examined by western blot analysis using anti-LGP2 antibody and anti-PC antibody.

To investigate the cause of the observed partial inhibition, we examined the knockdown efficiency of PC and LGP2 following siRNA treatment by western blot analysis. LGP2 siRNA treatment and si-PC treatment were able to reduce LGP2 and PC expression respectively (Fig. 6I-J). Given the limitations of PC being endogenous, mitochondrial and highly stable protein, we next employed PC knockout (KO) HEK cells to more precisely delineate the role of PC. PC-KO cells were reconstituted by transfection with their respective expression vectors (pPC-His), followed by transfecting miRTC14 to assess its functional effects. As a control experiment, an empty vector was used. Subsequently, IFN induction was quantified by employing qRT-PCR and ELISA assay. Overexpression of PC following pPC transfection was confirmed by WB analysis. As expected, PC was undetectable in PC-KO cells but was successfully restored upon transfection with the wild-type PC expression plasmid (pPC-wt) (Fig. 7A). Consistent with this, PC-KO cells failed to induce *IFNA2*, *IFNA8*, *IFNA13*, and *IFNL1* expressions in response to miRTC14 in both mRNA and protein levels (Fig. 7B-F). However, re-expression of PC rescued the miRTC14-induced upregulation of these IFNs, indicating that PC is essential for miRTC14-mediated interferon induction.

**Figure 7.**
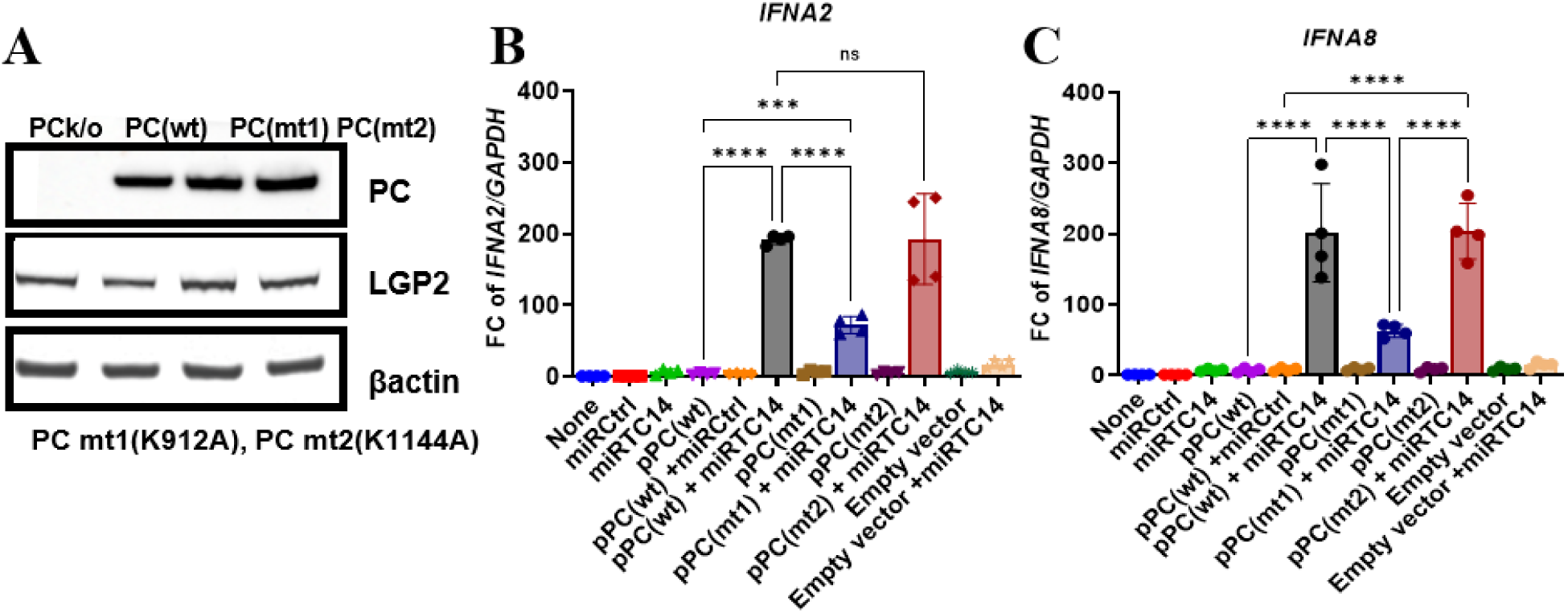
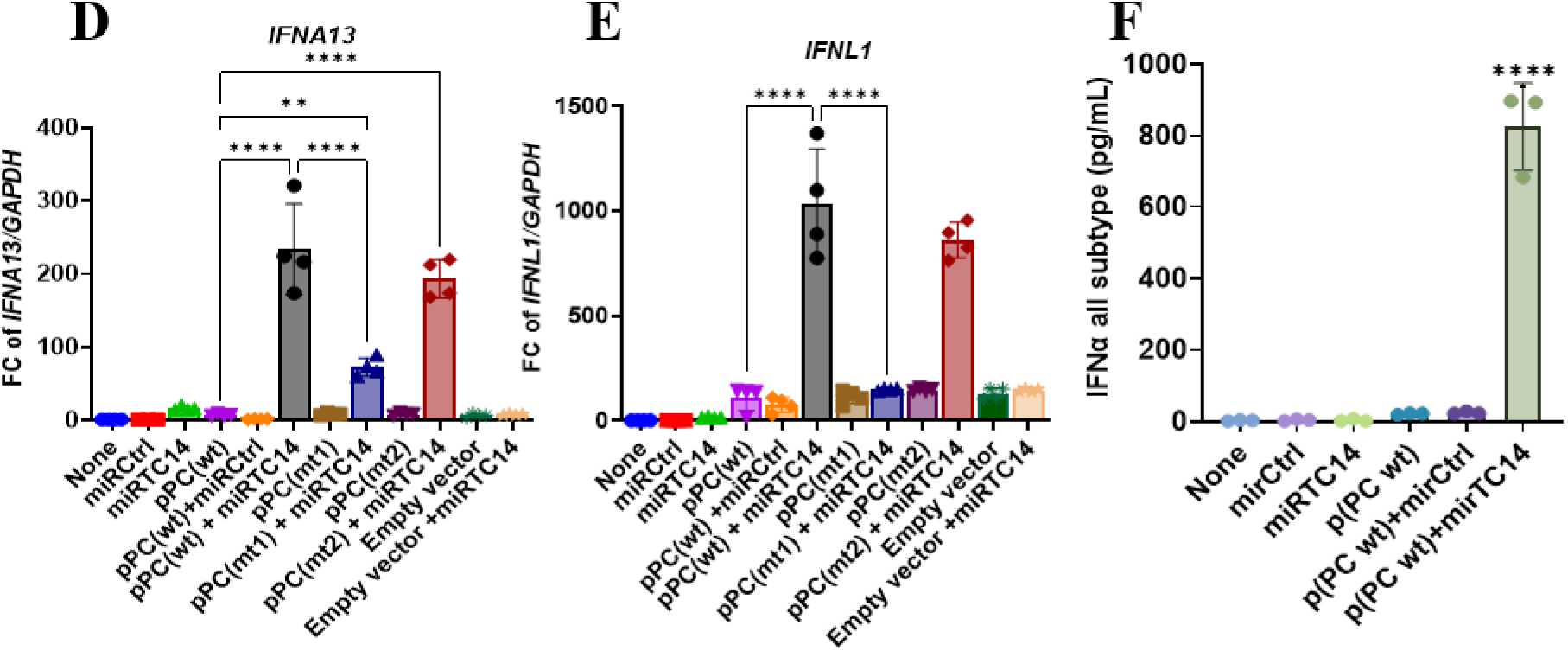
PC is required for miRTC14-mediated interferon induction. PC-KO cells were reconstituted by transfecting them with PC expression vectors (PC-wt, PC-mt1, PC-mt2). Expression of PC following plasmid transfection was confirmed by WB analysis **(A)**. miRTC14 and miRCtrl were transfected to PC overexpressed cells and lysed cells after 48 hrs of post-transfection. IFN expression was quantified by qRT–PCR. Gene expression is presented as relative expression units compared with no-transfection after normalization to GAPDH. Reconstitution of PC expression rescued miRTC14-induced upregulation of *IFNA2* **(B)**, *IFNA8* **(C)**, *IFNA13* **(D)**, and *IFNL1***(E)**, expression in response to miRTC14.Results show mean ± SD (*n* = 3). Secreted protein levels of IFN-α were measured using all subtypes IFN-α ELISA assay **(F)**. *, **, and *** indicate *p* values are < 0.05, < 0.01, and < 0.001, respectively.

### The binding pocket of PC is critical for the interaction between PC and miRTC14

To further characterize the interaction between PC and miRTC14 and to identify the potential binding domain/pocket and key residues in PC, we performed docking model analysis. Based on selected miRTC14-PC docking model, two hydrogen bonds were predicted to form between lysine residues at position 912 (K912) in PC and 2 phosphate groups between U_22_ and U_24_ (Fig. 5G-H). To further validate the predicted binding site, we performed the *in-silico* mutation experiments from PC. The results indicated that lysine amino acid (aa) at K912 was potential interacting site.

Based on these predictions, we generated point mutation in wild-type PC, namely K912A (PC-mt1) (Table S5). To validate the potential influence of biotinylation of the biotin-carboxyl carrier protein (BCCP) domain, we generated another point mutation, K1144A (PC-mt2). Mutant proteins were expressed and purified using a Ni-NTA column. Consistent with our previous experimental setup, PC-KO HEK cells were transfected with expression vectors encoding PC mt1 (pPC-mt1) or PC mt2 (pPC-mt2). Subsequently, miRTC14 was introduced to evaluate its functional effects. As a control, cells were transfected with an empty vector followed by miRTC14, and IFN induction was quantified by qRT-PCR. Overexpression of PC-mt1 and PC-mt2 in PC KO cells was confirmed by WB analysis (Fig. 7A). As described above, overexpression of wild-type PC (PC-wt) restored the ability of miRTC14 to induce *IFNA2, IFNA8, IFNA13*, and *IFNL1* expression (Fig. 7B-F). In contrast, mutation of K912A (PC-mt1) significantly attenuated miRTC14-mediated upregulation of these interferons (Fig. 7B-E), indicating that K912 is critical for miRTC14-induced IFN activation. Conversely, mutation of K1144A (PC-mt2) did not affect miRTC14-induced interferon expression compared to PC-wt (Fig. 7B-E). To further validate the effects of mutation of K912A and K1144A we performed binding assays using an Octet R8 instrument. Three forms of PC protein—PC-wt, PC-mt1 (K912A), and PC-mt2 (K1144A) were tested for interaction with miRTC14. Consistent with the IFN induction results, PC-mt1abolished binding to miRTC14, whereas PC-mt2 retained binding comparable to PC-wt (Fig. 8A), indicating that the BCCP domain undergoes a substantial conformational rearrangement remote from the binding site (Fig. 5F).

**Figure 8:**
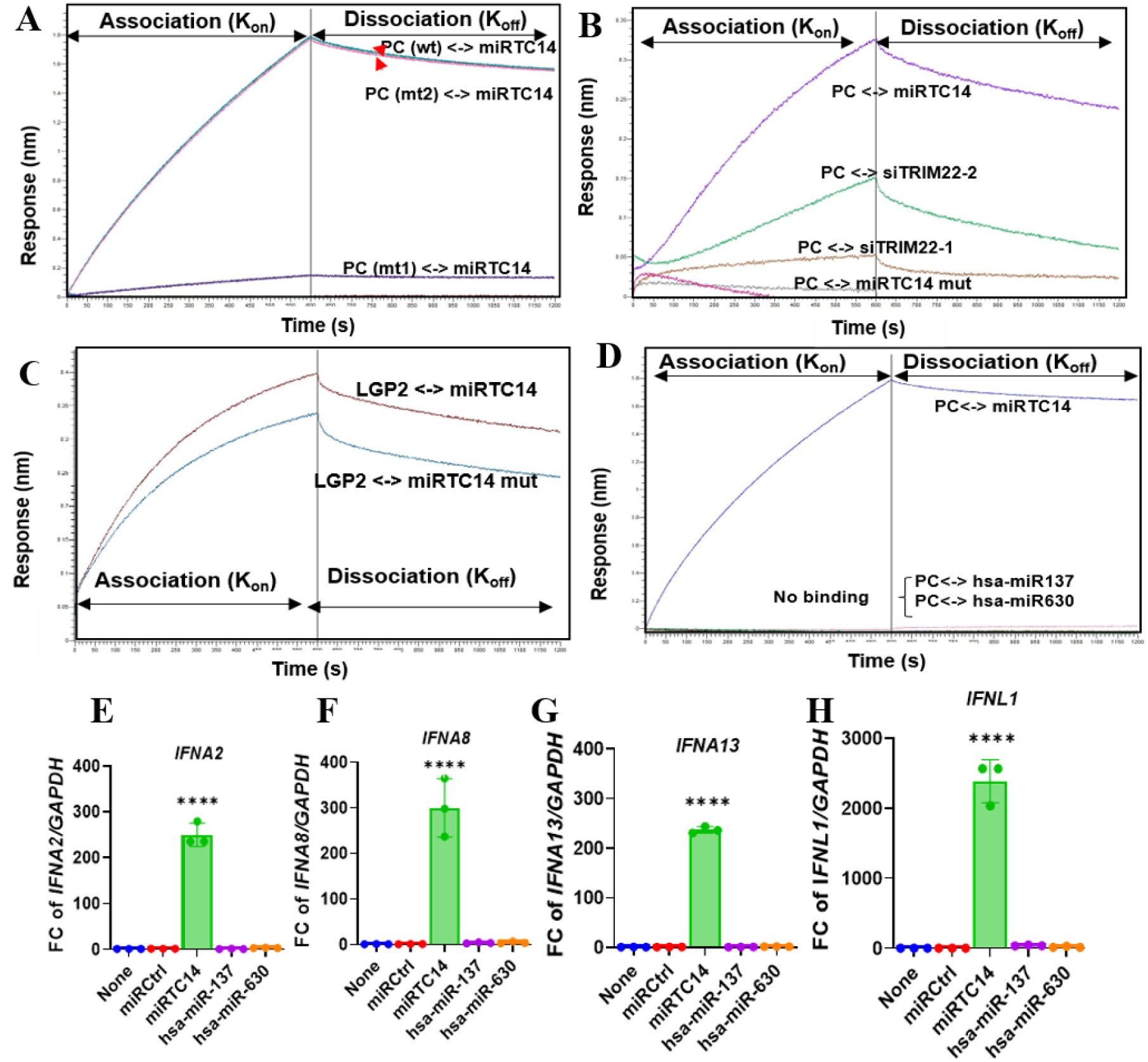
The binding pocket of PC mediates sequence-selective interaction with miRTC14. (**A**) Binding kinetics were measured using bio-layer interferometry. Recombinant PC proteins—wild-type (PC-wt), PC-mt1 (K912A), and PC-mt2 (K1144A)—were immobilized on Ni–NTA biosensors and assessed for interaction with miRTC14. The K912A mutation (PC-mt1) abolished binding, whereas K1144A (PC-mt2) retained binding comparable to PC-wt. (**B**) To evaluate binding specificity, purified 6×His-PC was immobilized on Ni–NTA sensors, and small RNAs (miRTC14-mimic, si-TRIM22-1, si-TRIM22-2, or miRTC14 mutants) were allowed to associate in solution. Mutant miRTC14 failed to bind PC but retained interaction with LGP2 (**C**), indicating distinct recognition requirements. (**D**) To evaluate binding ability to AUUCU-3′ sequence purified recombinant 6×His-PC proteins were immobilized on Ni–NTA sensor, while miRTC14, hsa-miR-137-5p and hsa-miR-630, were allowed to freely associate in solution. For all binding experiments, control measurements were performed in buffer alone. Kinetic parameters were derived by global fitting to a 1:1 binding model using the instrument analysis software, yielding association (k_on_) and dissociation (k_off_) rates. Representative sensorgrams are shown as solid lines (**E-H**). Sequence specificity of PC-mediated signalling was assessed in MDMs transfected with miRTC14 mimic, hsa-miR-137-5p, or hsa-miR-630. At 48 h post-transfection, expression of IFNA2 (**E**), IFNA8 (**F**), IFNA13 (**G**), and IFNL1 (**H**) was quantified by qRT–PCR. Gene expression was normalized to GAPDH and presented relative to non-transfected controls. Data represent mean fold change (FC) ± SD. (n=3). Statistical significance is indicated as *P < 0.05, **P < 0.01, ***P < 0.001.

### The interaction between miRTC14 and PC is both sequence-specific and site-specific

Based on the selected miRTC14–PC docking model, several hydrogen bonds were predicted between PC residue K912 and the 5’-U_22_C_23_U_24_-3’ region of miRTC14 along with additional non-bonded interactions (Fig. 5G-H). Based on *in-silico* mutation analysis, we found that most binding poses from the mutated miRTC14 (i.e., 5’-U_21_U_22_C_23_U_24_-3’ changed to 5’-A_21_G_22_A_23_A_24_-3’) exhibited suboptimal alignment with those of the wild-type miRTC14. This disruption indicates that these nucleotides are essential for maintaining the proper binding orientation and overall interaction stability with PC. Next, we performed binding assays using miRTC14, two siRNAs with partially similar sequences to miRTC14 (si-TRIM22-1 and si-TRIM22-2), and a mutated miRTC14 (Table 1). These assays were performed to evaluate the binding affinity of these miRNAs to PC protein. Consistent with previous observations, miRTC14 exhibited strong binding to PC, whereas the mutated miRTC14 lost this binding capability to PC (Fig. 8B). Despite the presence of comparable 5′-UUCU-3′ sequence motif, the two siRNAs exhibited markedly different binding behaviors. Among them, si-TRIM22-2 showed detectable interaction with PC, while si-TRIM22-1 exhibited no binding (Fig. 8B). However, the interaction between si-TRIM22-2 and PC was notably weaker compared to that of miRTC14 (Supplementary Table S3). A salient distinction between miRTC14 and si-TRIM22-2 is the absence of an additional adenosine residue preceding the 5′-UUCU-3′ motif in si-TRIM22-2 (Table 1). Together, this observation suggests that the presence of an upstream adenosine in the motif may contribute to efficient interaction with PC, and that the 5′-AUUCU-3’ (and corresponding 5′-AGAAU-3′) sequence context is critical for binding and may be for downstream IFN induction. However, the mutated miRTC14 only modestly affected its binding to LGP2 (Fig. 8C), suggesting that the PC and LGP2 binding sequences are distinct.

To further confirm the site specificity, we examined additional miRNAs containing the 5′-AUUCU-3′ motif. Two such candidates, hsa-miR-137-5p and hsa-miR-630, were identified from publicly available databases (Table 1). Binding assays were then performed using miRCtrl, miRTC14, hsa-miR-137-5p, and hsa-miR-630 in the presence of PC. As expected, only miRTC14 exhibited binding to PC, whereas the other microRNAs showed no detectable interaction (Fig. 8D), indicating that the presence of the 5′-AUUCU-3′ motif alone is insufficient and that site specificity is critical for PC binding. To assess functional consequences, each miRNA was transfected into MDMs, followed by qRT-PCR analysis of IFN gene expression. Consistent with the binding results, neither hsa-miR-137-5p nor hsa-miR-630 induced IFN expression (Fig. 8E-H). Taken together, these findings demonstrate that both sequence composition and positional context are required for efficient PC recognition and subsequent IFN induction by miRTC14.

Based on the above findings, we now know that both PC and LGP2 are required for the miRTC14-mediated IFN induction. To investigate the interaction dynamics among PC, miRTC14, and LGP2, we analyzed their binding behavior over time using Octet R8. First, we allowed PC to bind miRTC14, followed by the addition of recombinant LGP2. The binding data showed sequential engagement of miRTC14 with the two proteins (Supplementary Fig. S10A). In parallel, we also tested the reciprocal condition by introducing PC to preformed LGP2–miRTC14 complexes. However, when we introduced PC to LGP2–miRTC14 complexes, the binding was relatively low, suggesting differences in binding stability, conformation, or detection efficiency between the PC-bound and LGP2-bound states (Supplementary Fig. S10B). These differences demonstrate that, although both PC and LGP2 bind the same miRTC14, they recognize and engage distinct RNA regions or conformational state of miRTC14.

### PC acts as a selective RNA sensor, with no detectable response to DNA in triggering interferon signalling

Since we identified PC as a selective RNA sensor/ binding protein and next investigated its potential role in DNA sensing. Given that sequence and site specificity are critical for miRTC14–PC–mediated IFN induction, we designed a double-stranded DNA construct (dsDNA14), based on a sequence analogous to miRTC14. To assess IFN inducibility, dsDNA14 was transfected into MDMs and IFN expression was quantified by qRT-PCR. Despite its sequence similarity to miRTC14, dsDNA14 did not induce IFN expression, indicating that PC selectively recognizes RNA rather than DNA in triggering IFN signalling (Supplementary Fig. S11). The inability of dsDNA14 to induce IFN reinforces the concept that PC functions as a selective RNA sensor, likely recognizing features such as RNA backbone chemistry, duplex dynamics, or end structures that are critical for downstream activation.

### LGP2 is critical for miRTC14-mediated interferon induction

Similar to PC-KO experiment in Fig. 7, LGP2-KO cells were transfected with LGP2 expression vectors (pLGP2-His), and empty vector as a control experiment. Then miRTC14 was transfected. Subsequently, IFN induction was quantified by employing qRT-PCR and ELISA assay. LGP2 was absent in LGP2-KO cells; although, PC was endogenously expressed in these cells. LGP2 expression restored upon transfection with pLGP2 (Fig. 9A). In LGP2-KO cells, even the presence of PC, miRTC14-induced expression of *IFNA2*, *IFNA8*, *IFNA13*, and *IFNL1* were markedly impaired. Importantly, reconstitution of LGP2 expression restored the ability of miRTC14 to induce these interferons (Fig. 9B-E). This finding was further confirmed by ELISA assay by quantifying total IFNα subtype protein levels (Fig. 9F). Taken together, these findings demonstrate that both PC and LGP2 are required for miRTC14-mediated IFN induction, suggesting that this response is driven by a cooperative mechanism involving both proteins.

**Figure 9.**
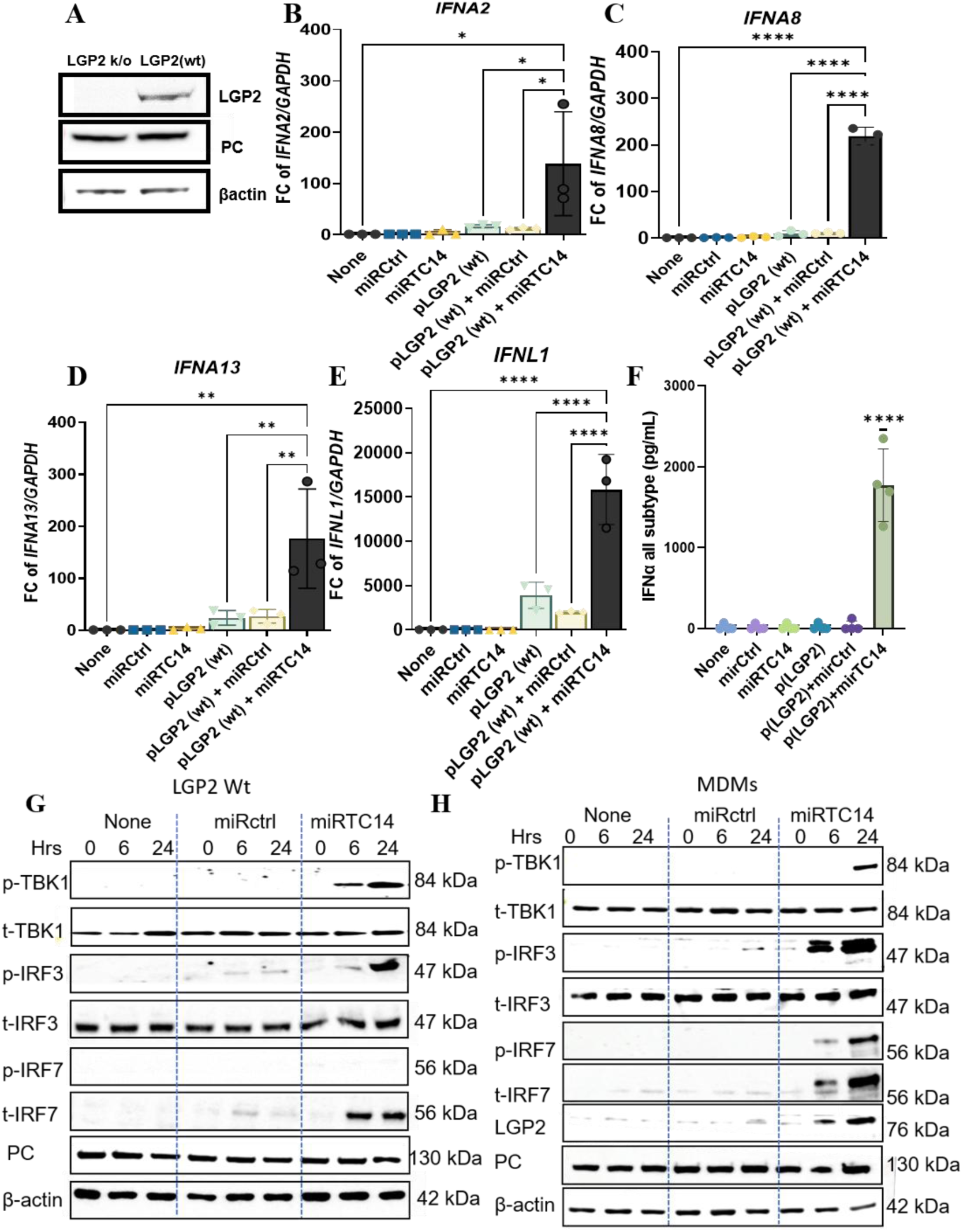
LGP2 is critical for miRTC14-mediated interferon induction. LGP2-KO cells were reconstituted by transfecting with wild-type LGP2 expression vectors (LGP2-wt). Overexpression of LGP2 following plasmid transfection as well as the expression of endogenous PC were confirmed by WB analysis **(A)**. miRTC14 and miRCtrl were transfected to LGP2 overexpressed cells and lysed cells after 48 hrs of post-transfection. IFN expression was quantified by qRT–PCR. Gene expression is presented as relative expression units compared with no-transfection after normalization to GAPDH. Reconstitution of LGP2 expression rescued miRTC14-induced upregulation of *IFNA2* **(B)**, *IFNA8* **(C)**, *IFNA13* **(D)**, and *IFNL1***(E)**, expression in response to miRTC14. Results show mean ± SD (*n* =3). Secreted protein levels of IFN-α all subtypes were measured using ELISA assay **(F)**. *, **, and *** indicate *p* values are < 0.05, < 0.01, and < 0.001, respectively. (**G**) LGP2-KO cells were reconstituted by transfecting LGP2 expression vectors (LGP2-wt) and cultured for 72 hours. LGP2-wt transfected cells (**G**) and MDMs (**H**) were then transfected with miRTC14 and miRCtrl and lysed cells after 0, 6 and 24 hours of post-transfection, and western blotting was performed using anti-p-TBK1, anti-t-TBK1, anti-p-IRF3, anti-t-IRF3, anti-p-IRF7, anti-t-IRF7, LGP2 and PC antibodies. Anti-β-actin antibody was used to probe β-actin as a loading control for all the samples.

### PC-miRTC14-LGP2-axis induces TBK1 phosphorylation

LGP2 has been shown to promote activation of TANK-binding kinase 1 (TBK1), thereby facilitating dsRNA-mediated canonical NF-κB and IRF3 signalling pathways (66). Given our observation that miRTC14-PC complex binds to LGP2 and induces IFN expression, we next examined TBK1 phosphorylation and its downstream signalling cascade in miRTC14–transfected cells. To assess the specific involvement of LGP2, we utilized both LGP2 KO cells and LGP2-overexpressing cells, analysing responses at 6 and 24 hours post–miRTC-14 transfection. WB was performed to test the phosphorylation of TBK1, IRF3 and IRF7. Total TBK-1, IRF3 and IRF7 were used as an internal references and β-Actin was used as a loading control. LGP2 knockout cells failed to exhibit TBK1 phosphorylation in response to miRTC14 (Supplementary Fig. S12). In contrast, LGP2 overexpression induced robust TBK1 phosphorylation at both 6 and 24 hours following miRTC14 transfection, but not with miRCtrl (Fig. 9G). A similar phosphorylation pattern was observed for phospho-IRF3, whereas phosphorylated IRF7 was not detected (Fig. 9G). In MDMs, however, miRTC14 transfection increased phosphorylation of TBK1, IRF3, and IRF7, indicating the involvement of additional regulatory steps compared with HEK cells (Fig. 9H). Notably, phosphorylation intensity was more pronounced at 24 hours post-transfection, suggesting a potential sequential amplification mechanism underlying miRTC14-mediated innate immune responses.

## DISCUSSION

In this study, we found a non-canonical mechanism by which an IL-27–induced microRNA, miRTC14, promotes antiviral immunity through the induction of type I and type III IFNs. We demonstrated that a previously unrecognized axis involving the metabolic enzyme PC and the RNA sensor LGP2/DHX58. These findings established a novel RNA-sensing mechanism that links cellular metabolism to innate immune activation.

A striking feature of miRTC14-PC-LGP2 axis is its selective induction of IFN genes. In humans, 13 distinct IFN-α genes have been identified (49,50). In addition, one copy of the IFN-β, IFN-Epsilon, IFN-Kappa, and IFN-Omega genes has been documented (49). Although human IFN-α genes are clustered on chromosome 9, and IFN-β is also mapped to the same chromosome, the axis activated IFNA2, A4/10, A5, A8, and B genes by more than 100-fold and induced IFNA1, A13, A14, and A17 genes by more than 40-fold. It is noteworthy that IFN-A6 and IFN-A16 genes were not activated. This selective responsiveness indicates that IFN loci are differentially poised for activation downstream of miRTC14 signalling. Such specificity may reflect promoter accessibility, chromatin context, or transcription factor recruitment, and is consistent with emerging evidence that IFN gene activation is highly modular rather than uniform.

A key finding of this work is the identification of PC as a direct binding partner of miRTC14, alongside the RNA sensor LGP2. While LGP2, a well-characterized RNA sensor, has been demonstrated to play a crucial role in the process of MDA5-mediated innate immunity (65,67,68), the identification of PC as an RNA-binding protein is unexpected, as it is classically defined as a mitochondrial metabolic enzyme with unknown role in RNA recognition. However, our combined biochemical and computational analyses which include pull-down assays, biolayer interferometry, confocal imaging, and *in silico* docking, demonstrate that miRTC14 binds with PC with relatively high affinity and adopts a consistent conformation within a predicted C-terminal pocket. These findings suggest that PC possesses previously unrecognized RNA-binding capacity, expanding its functional repertoire beyond metabolism. Kinetic measurements further indicate that the stronger miRTC14–PC interaction than the miRTC14–LGP2, implying that LGP2 may play a dominant role in stabilizing signalling complexes that drive IFN induction. In this model, PC may function as an initial RNA docking scaffold or conformational regulator that facilitates subsequent engagement with LGP2. The subcellular localization of this pathway presents an additional layer of complexity (14). Although PC is predominantly a mitochondrial matrix enzyme that forms tetramers (49) and LGP2 is a cytosolic protein (65), confocal imaging revealed miRTC14 localization within mitochondria, challenging the assumption that complex formation occurs in the cytosol prior to mitochondrial import and instead raises the possibility that RNA sensing may occur in proximity to, or within, mitochondrial-associated microenvironments. Previous work has been shown that upon viral infection, RIG-I activation displaces LGP2, enabling innate immune signalling and IRF3/7-driven interferon responses (14). In the current study, we demonstrated that the PC–miRTC14–LGP2 axis induced TBK1 activation in a similar manner to other RNA-mediated innate immune responses, such as those triggered by RIG-1 or MDA5 (12,65). Although TBK1 was activated, the extent of gene induction differed among IFN subtypes, indicating the involvement of additional regulatory mechanisms. These may include differential recruitment of transcriptional co-activators, promoter architecture, or feedback mechanisms that fine-tune IFN responses. Following complex formation, PC–miRTC14 undergoes structural destabilization, potentially exposing the LGP2 binding site and leading to TBK1 activation.

Collectively, our findings expand the conceptual framework of microRNA function by demonstrating that specific RNA sequences can act as immunostimulatory ligands. This study highlights the importance of sequence context and higher-order RNA features in determining protein recognition and downstream signalling outcomes. The identification of PC as an RNA-binding protein further suggests that metabolic enzymes may serve as previously unappreciated components of innate immune sensing pathways, thereby providing a mechanistic link between metabolism and antiviral defence. Since canonical innate immune response against RNA causes sequence independent IFN gene activation, preferentially induces IFN-β more than IFN-α, this PC-mediated selected IFN-α subset gene activation is non-canonical innate immune response.

Beyond its established function in TCA cycle anaplerosis and roles in cellular senescence, PC emerges here as a moonlighting protein linking metabolism to antiviral immunity. From a translational standpoint, miRTC14 or engineered mimetics may represent promising candidates for antiviral therapy, particularly against HIV and IFN susceptible viruses. More broadly, the ability to design small RNAs that selectively engage immune pathways through defined sequence and structural features offers a potential strategy for immunomodulation. Future studies will be required to resolve the high-resolution structure of the miRTC14–PC complex and to determine how this pathway interfaces with established RNA sensors. Such insights may reveal new opportunities for targeting antiviral intervention.

## INSTITUTIONAL REVIEW BOARD STATEMENT

Approval of these studies, including all sample materials, was granted by the National Institute of Allergy and Infectious Diseases Institutional Review Board. All experimental procedures in these studies were approved by the National Cancer Institute at Frederick and Frederick National Laboratory for Cancer Research and were performed in accordance with the relevant guidelines and regulations (the protocol code number: IBC: 2016-19 A8/11-30, approval data: 8 August 2023).

## ACKNOWLEDGEMENTS

The authors thank H.C. Lane for discussing this project, G. Panzade for critical reading and H.-N. Sharm for analysis of database. The content of this publication does not necessarily reflect the views or policies of the Department of Health and Human Services, nor does mentioning trade names, commercial products, or organizations imply endorsement by the U.S. Government. This research was supported (in part) by the National Institute of Allergy and Infectious Diseases. The pHIV_AD8_ and pHIVNL4-3 ΔEnv/Vpr Luciferase were obtained through BEI Resources, NIAID, NIH.

## AUTHOR CONTRIBUTIONS

Conceptualization: T.I., U.K., and S.G. Methodology, data analysis, and interpretation: U.K., S.G., RW.M., Q.C., J.Y., M.M., H.S., and T.I. Visualization: U.K., and T.I. Writing – original draft: U.K., T.I., M.H., and H.S. Writing – review and editing: U.K., S.G., M.H., RW.R., Q.C., J.Y., J.Q., M.M., H.S., W.C. and T.I. Bioinformatic analyses: M.H., and W.C. Supervision: T.I. Funding acquisition: T.I.

## SUPPLEMENTARY DATA

Supplementary Data are available at NAR online.

## DATA AVAILABILITY

All data needed to evaluate and reproduce the results in the paper are present in the paper and/or the Supplementary Materials.

