## Supplemental Table 2 for "Beyond Metabolism: Pyruvate Carboxylase Acts as a Sequence-Selective Small RNA Sensor Orchestrating Antiviral Responses"

**Supplementary Table S2: Gene names of predicted using Venn Diagram**

| Names | total | elements |
|---|---|---|
| miRAB40 miRTC10 miRTC14 | 2 | ATP5F1E |
| | | KCMF1 |
| SX2 miRAB40 miRTC10 | 8 | ROCK2 |
| | | TBL1XR1 |
| | | SMC1A |
| | | APPL1 |
| | | SLC16A10 |
| | | RASSF8 |
| | | ZNF704 |
| | | HIPK1 |
| SX2 miRAB40 miRTC14 | 1 | MTREX |
| SX2 miRTC10 miRTC14 | 11 | ABCC5 |
| | | PARD6B |
| | | ENAH |
| | | TMEM242 |
| | | SETX |
| | | SLC35B4 |
| | | ZNF785 |
| | | GM2A |
| | | SLC25A33 |
| | | STK35 |
| | | CYTH3 |
| miRAB40 miRTC10 | 39 | KBTBD8 |
| | | GDA |
| | | HSPD1 |
| | | CIP2A |
| | | SKAP2 |
| | | TMOD2 |
| | | NR1D2 |
| | | FMR1 |
| | | KCNG3 |
| | | ZNF550 |
| | | ITCH |
| | | PPP1R21 |
| | | SFT2D2 |
| | | NXPE3 |
| | | ZNF697 |
| | | PIK3C3 |
| | | TMEM47 |
| | | SENP6 |
| | | CPM |
| | | C8orf34 |
| | | PRICKLE2 |
| | | CSRNP3 |
| | | SNX27 |
| | | DCUN1D4 |
| | | QSER1 |
| | | DTWD1 |
| | | BMT2 |
| | | ABI2 |
| | | OSBPL10 |
| | | CFAP44 |
| | | BARD1 |
| | | IMPACT |
| | | PMAIP1 |
| | | RAB9A |
| | | FBXO28 |
| | | PTAFR |
| | | GXYLT1 |
| | | HEY2 |
| | | BCKDHB |
| miRAB40 miRTC14 | 8 | RPRD1B |
| | | PAN3 |

| | | |
|---|---|---|
| | | TBL1X |
| | | ANP32E |
| | | C1orf131 |
| | | ZNF596 |
| | | HTD2; RPP14 |
| | | CYP20A1 |
| SX2 miRAB40 | 15 | MOB1B |
| | | TMBIM1 |
| | | RSF1 |
| | | PTAR1 |
| | | HIBCH |
| | | IER3IP1 |
| | | RADX |
| | | TOR1AIP1 |
| | | ZNF365 |
| | | BHLHE41 |
| | | RASSF3 |
| | | NRAS |
| | | UBE2D1 |
| | | TNFSF13B |
| | | UBE3D |
| miRTC10 miRTC14 | 227 | MON1B |
| | | PDLIM2 |
| | | TP53AIP1 |
| | | STK25 |
| | | GADL1 |
| | | USP6NL |
| | | HS2ST1 |
| | | CPEB3 |
| | | MYORG |
| | | SH3PXD2A |
| | | SPINK5 |
| | | OPA3 |
| | | IL17D |
| | | ITSN1 |
| | | PSTK |
| | | ATP2B2 |
| | | KLHL21 |
| | | TANC2 |
| | | NAB2 |
| | | POLR1A |
| | | NF1 |
| | | CCDC144A |
| | | NFAT5 |
| | | ARHGAP26 |
| | | CDS2 |
| | | POP4 |
| | | GALNT10 |
| | | RHBG |
| | | EFR3B |
| | | TRIM24 |
| | | SLC22A5 |
| | | GRAMD1B |
| | | OCEL1 |
| | | GFOD1 |
| | | ABCB9 |
| | | DHX38 |
| | | RGS11 |
| | | APOLD1 |
| | | CMTM4 |
| | | MCM2 |
| | | MTPN |
| | | SNX22 |
| | | PDE4DIP |
| | | NADK |
| | | NEURL4 |

NREP
MASP1
RIMKLA
MBOAT7
ZNF814
DEXI
ABCA5
WFDC6
MCPH1
SFXN5
ESPL1
ERGIC1
KIF26B
TVP23C
WWC2
AC009133.6
KIAA1522
NOS1
INTU
RBM24
CNKSR3
FAT2
ISL2
RBMS2
RBM19
CCDC32
PPP2R5C
ST3GAL6
ARSB
CLMN
C4orf33
GNL1
GREM1
SULT4A1
SLC28A1
VPS35L
AGO1
DFFA
ELP5
DCUN1D1
OGFOD3
WDR35
STRBP
AC009163.3
C19orf54
SERAC1
C2CD2
FAM3A
BAZ1A
MYO1F
CHD5
SETD7
KIAA1328
HIPK2
LILRB1-AS1
USP22
GPRIN3
RNF130
WNT4
RASGRF2
TMEM101
NF2
EPB41L5
E2F3
SSTR5
FOSL1

ATRN
AMZ1
SLC7A6
ITPK1
NKX2-5
PCGF3
ITM2B
BSN
CECR2
SPIRE2
DAPK3
BTN3A1
AC020929.1
CBFA2T2
DISC1
CLTCL1
LRRC2
CSPP1
CYB5R4
SCN4B
CRCP
TOGARAM2
ATXN2
TEX14
DRP2
TSPAN31
GFRA2
ALS2
NME9
ADARB1
SYNE3
TSC1
NARF
FTO
UBE3B
NKAIN4
BTN2A2
TMED8
MYLK
SNRK
EHMT1
SELENOI
ADGRF3
FGD3
MRPS25
ELMSAN1
MRPL42
PAGR1
AC006254.1
TRIM36
TMC7
XYLT1
AGO3
SYAP1
CCDC25
RGMB
PWWP3A
PLCB2
SSH2
ZNF500
GABPB2
KLF16
SWAP70
USP49
CLPTM1
ZNF707

| | | |
|---|---|---|
| | | GNB1L |
| | | PARP14 |
| | | MYT1 |
| | | DLX6 |
| | | ARL10 |
| | | PTK7 |
| | | B4GALT2 |
| | | RAB6B |
| | | FAM168B |
| | | DMRTA1 |
| | | DIRAS2 |
| | | AMDHD2 |
| | | TJP1 |
| | | TRERF1 |
| | | TADA2A |
| | | COX19 |
| | | SH3TC1 |
| | | ILF3 |
| | | CCND3 |
| | | FOXK1 |
| | | ETF1 |
| | | CREB5 |
| | | SHF |
| | | DACH2 |
| | | PPARA |
| | | TPM4 |
| | | CEP78 |
| | | MTG2 |
| | | ATAD2 |
| | | NLRC5 |
| | | CCDC191 |
| | | KREMEN1 |
| | | LIFR |
| | | CBL |
| | | SLC18A2 |
| | | EPG5 |
| | | PTS |
| | | MED28 |
| | | HBS1L |
| | | CCDC150 |
| | | SIK3 |
| | | KSR2 |
| | | BMPR1B |
| | | RNF157 |
| | | PEX5L |
| | | PRR26 |
| | | TTC9 |
| | | LRRC14 |
| | | PRMT7 |
| | | CTSC |
| SX2 miRTC10 | 188 | SCAF8 |
| | | C2orf49 |
| | | DDX5 |
| | | BORCS7 |
| | | RFXAP |
| | | EIF1AX |
| | | EXOC2 |
| | | L2HGDH |
| | | KLHL7 |
| | | DCUN1D5 |
| | | GMFB |
| | | NFYA |
| | | TLK1 |
| | | CPD |
| | | LAMTOR3 |
| | | ZBTB8OS |

MKLN1
TRHDE
ZNF677
ITGA2
C6orf120
ATG3
ST6GALNAC3
USP38
SEC22C
LNPK
TRAK2
ZNF544
TRIM11
RAD50
KANSL3
ZNF688
KIF3A
PIK3R1
C22orf39
CAV2
NIPAL1
CYB5A
SLIT2
LYRM7
DTNA
PSPH
C5orf24
COG6
RBBP9
RBM33
CHEK1
GNG10
ALG14
ZNF641
CYP4F3
ZNF366
PSMC6
DYRK2
DYNLL2
DBT
SCD
CACUL1
GID4
MPZL1
SHPRH
TNKS
LTN1
POLR3E
ZSWIM6
AMIGO1
NUDT3
RAP2B
CREB3L2
USP31
ZNF766
TDG
FIGNL1
ZNF773
RNF144B
EHD3
BAG2
SHISA9
SPRY3
SURF6
MRS2
MSRA

RBM18
NOL9
NIT2
CBX5
ZNF37A
EOGT
STUM
IL7
DPYSL3
CIPC
PARP16
MTRR
IKZF3
HINT3
SLC35A5
TENT5C
WBP11
IMPA2
TPD52L1
BLOC1S5
PIGK
SSBP2
REEP1
TRIP11
DIAPH3
PRR13
SMAD2
TLN2
TBCEL
MAPK1
USP46
PGAP1
SLC35D1
RNF24
IFNLR1
PCDH17
CLIP4
ZNF772
TGFBR3
PPA2
RABL2B
DNAL1
RAB18
RBM28
NUDT21
ARL8B
COMT
TMTC2
PGK1
DNAJB1
CCDC50
MAN2A2
ACOX1
C1orf56
NECAP2
CAVIN1
DHX40
TTBK2
FBXW2
HDAC9
RAB21
UHMK1
AGO4
LEPROT
KCNMB1
POGLUT3

| | | |
|---|---|---|
| | | AP5M1 |
| | | ATP5F1A |
| | | MTERF4 |
| | | PPIA |
| | | NXF1 |
| | | AK2 |
| | | SNX20 |
| | | GOPC |
| | | NIPAL2 |
| | | LONP2 |
| | | ZMAT3 |
| | | TRAPPC13 |
| | | KIAA0040 |
| | | GPR82 |
| | | ZBTB42 |
| | | KDR |
| | | LYRM9 |
| | | INHBA |
| | | RAP1B |
| | | PLXDC2 |
| | | CYGB |
| | | GALK2 |
| | | GNAQ |
| | | TMEM245 |
| | | TNFAIP8 |
| | | TNIP2 |
| | | SORD |
| | | TGFA |
| | | DR1 |
| | | ZNF180 |
| | | EIF4E3 |
| | | AAGAB |
| | | FYTTD1 |
| | | PDK1 |
| | | ZNF736 |
| | | LRRC28 |
| | | DUSP16 |
| | | LARP4B |
| | | CAST |
| | | UBE2D2 |
| SX2 miRTC14 | 45 | SRPK1 |
| | | CCDC34 |
| | | PAQR8 |
| | | NEGR1 |
| | | SC5D |
| | | WSB1 |
| | | ZBTB7C |
| | | RMND5A |
| | | SNX18 |
| | | ABHD8 |
| | | FZD3 |
| | | DMWD |
| | | ZNF689 |
| | | RPS23 |
| | | SMAD3 |
| | | NAPG |
| | | FAR2 |
| | | ARIH2 |
| | | UQCRB |
| | | XPR1 |
| | | NDUFV3 |
| | | CYREN |
| | | USP47 |
| | | STMP1 |
| | | LCMT2 |
| | | YWHAG |

| | | |
|---|---|---|
| | | SRSF7 |
| | | RHOU |
| | | DAZAP2 |
| | | LUC7L3 |
| | | FSBP |
| | | EIF4H |
| | | CALM1 |
| | | AIDA |
| | | MAP2K6 |
| | | KXD1 |
| | | RAN |
| | | PHF6 |
| | | MXI1 |
| | | TCAF2 |
| | | CALU |
| | | CALCRL |
| | | DEPTOR |
| | | FAM110B |
| | | SLC38A7 |
| miRAB40 | 115 | TPM3 |
| | | SPZ1 |
| | | TAF12 |
| | | RGS4 |
| | | DAPP1 |
| | | ANKLE2 |
| | | OSBP |
| | | ZNF667 |
| | | RAB15 |
| | | B2M |
| | | TRPC1 |
| | | OSTN |
| | | H2AZ1 |
| | | TECTB |
| | | C9orf153 |
| | | RFPL3 |
| | | TUT4 |
| | | NLK |
| | | ITGBL1 |
| | | RASGEF1B |
| | | ZNF624 |
| | | PRR20B |
| | | ZBED9 |
| | | SLC24A1 |
| | | PRR20A |
| | | SLC39A10 |
| | | KCNV1 |
| | | NCBP3 |
| | | SLC46A3 |
| | | MPEG1 |
| | | DMD |
| | | CHURC1 |
| | | CCP110 |
| | | NOXRED1 |
| | | CLGN |
| | | CDK15 |
| | | YIPF4 |
| | | ETS2 |
| | | FCRL3 |
| | | RNF144A |
| | | RNF103 |
| | | ZIC4 |
| | | RRP36 |
| | | ITGB6 |
| | | ZSCAN29 |
| | | SETD5 |
| | | GABRA2 |

DENND5B
CAMK2N1
TOPORS
TMEM150C
CUL5
TPGS2
HTATIP2
NMNAT3
FAM241A
GIN1
GSTM3
KIF24
CSNK1G3
DTHD1
SMIM15
CLLU1OS
MAP4K4
LMNA
BTG3
RTKN2
CRTAM
NT5DC1
CAND1
SPTSSA
TRA2B
OTUD6B
SMAD9
TMEM87B
DHX15
SEC63
ITGB1
TEX2
BROX
PRR20E
ZFX
HAS3
HLA-DPB1
GPR85
MAP3K7
EID2
SLC7A1
SIK1
PCDHA6
OCLM
ARHGAP11A
GOLM1
USP24
PINLYP
POU5F2
CLVS2
USP25
TPM1
PRPF38B
RPRD2
KCNJ1
PRR20D
FGF7
ZNF286A
KIAA1671
FNDC3A
BICC1
SCAMP1
SERTM1
MGAT4A
ZNF286B
PRDM15

| | | |
|---|---|---|
| | | FGF23 |
| | | RFPL1 |
| miRTC10 | 2782 | SLMAP |
| | | SAMD4A |
| | | FAM219A |
| | | C10orf25 |
| | | H2AZ2 |
| | | PDCL3 |
| | | ZNF646 |
| | | RAB1B |
| | | HOMEZ |
| | | TOP1MT |
| | | CRNKL1 |
| | | C12orf66 |
| | | SLC26A1 |
| | | HROB |
| | | NUTM2B |
| | | DIRAS1 |
| | | CCDC28B |
| | | KCNK3 |
| | | MSRB3 |
| | | COX15 |
| | | PDE1C |
| | | SEMA4D |
| | | TOB2 |
| | | INIP |
| | | MAP3K3 |
| | | TBC1D3H |
| | | RRP7A |
| | | TMEM132E |
| | | PTCD3 |
| | | SORL1 |
| | | MAPKAPK2 |
| | | BRCA1 |
| | | SLK |
| | | PRKCB |
| | | EIF4G3 |
| | | NOL8 |
| | | CDK20 |
| | | RELT |
| | | CCDC66 |
| | | BBS12 |
| | | LARP7 |
| | | TERF2IP |
| | | PINX1; PINX1; SOX7 |
| | | NUTM2G |
| | | PHF20 |
| | | HNF1A-AS1 |
| | | PDCD5 |
| | | HES1 |
| | | TMEM52 |
| | | LINC01620 |
| | | GAGE1 |
| | | ZNF212 |
| | | GRK1 |
| | | VGLL3 |
| | | PRDM2 |
| | | ZNF83 |
| | | SEL1L3 |
| | | SUPT4H1 |
| | | SRRM4 |
| | | RNF223 |
| | | GFPT2 |
| | | KCNC1 |
| | | ARHGEF9 |
| | | OR8J1 |

CCL19
GPR55
GBP7
STAU2
VWA5A
ABR
PGBD5
CTDSP2
DUSP4
ZNF257
GPR78
ISM1
CORO2A
VAT1L
PSMG4
C9orf139
ING5
ALG2
PPP1R3D
GRB2
ATXN7L3B
OR4A47
SYT13
ORM1
PIWIL3
ZNF232
DESI1
PANX3
PLXNA1
C8orf44-SGK3
MAP3K13
LCMT1
PPM1A
BCL2
MMP16
ARL13B
TJAP1
PGGHG
AIPL1
SNX19
THADA
PIK3CB
PDZD3
FAXC
FAM110C
TOMM34
CD6
CXorf56
DNAL4
PSMD11
ELAC2
ALAD
USP42
CDC42BPA
AC137056.1
B3GALT5
TFR2
IRS1
TMEM104
MFN2
PTPN5
ZSCAN32
OGFOD2
DDX49
FAM204A
SEPTIN4

MYO5C
ZBED1
ANKRD49
POF1B
ZNF552
KDM2A
CACNB1
LDHAL6A
KDM6A
QKI
LAMB4
AHCYL2
KDM4B
TARS1
HABP4
CNTN2
IL6R
CHRNB2
UBN2
FUT4
DBH
CIITA
MUTYH
ZNF655
ZNF227
MCOLN2
CDH8
NEU1
PCCB
MAK16
EPGN
OSR2
LZTFL1
ARHGAP15
REM2
SMTNL2
LONRF2
GIPR
HCFC1
GORAB
DEFB125
UBE2W
KRT15
IL18BP
SCN11A
MYLK2
PTPRH
VSIR
POTEG; POTEM
ATP10A
CAMSAP1
XCR1
KCNA4
PALM2AKAP2
RBM6
NMNAT2
MAGEB6
IFITM10
MRPS24
SCHIP1
JADE1
DYRK4
UIMC1
TFEC
DCX
RIC3

H2BC18
POLRMT
ZAN
CRYBA1
EIF4E2
HRH2
SOX9
PDGFRB
FHDC1
GPR158
CARD9
RDX
PACSIN3
HOXB8
JUND
TMEM241
MRTO4
TMEM248
TCF21
SLA
STOML3
TCP1
TBC1D3B
ESPNL
FAM177A1
RER1
DDI2
GPATCH8
ZNF586
SNCAIP
ANXA8
DCT
KLHL28
FBF1
PELI3
SET
SHH
EIF2S1
HSFY2
GPR135
PIGG
AL354861.3
C1orf115
ZNF264
BICD1
DCAF11
ACADVL
GTF2H5
PLEKHM3
ICA1L
HCAR2
XIAP
SPAG6
ZDHHC22
RPS29
LGALS4
DAGLB
PPFIBP1
GLIPR2
TRIM63
PLRG1
YTHDC2
ZNF614
SCRT2
XXYLT1
ELF1

BTBD9
GK5
HHIPL1
NOTCH2
CD93
ZSCAN12
PSMD12
HSCB
GYPA
AP001267.2
MARCHF4
FKBP5
SMS
CHCHD5
SPDYE1
FBXO3
CMKLR1
CSNK2A2
SDK2
ANKRD40
GPX2
SPATS2L
ZNF551
BSDC1
PHF12
CTCFL
HS3ST5
TEKT2
PSMB8
SDR39U1
SETBP1
GIMAP6
MAST4
EEF2K
TRMT2B
KCTD2
SPTLC3
TMEM207
UNKL
MAPK8IP2
POU3F2
ALPK1
MCAT
ZFP1
VPS50
WDPCP
MAGI1
VCP
SLC35A1
RAVER1
TSTD1
GPD1L
MOSMO
ADAMTS17
PANK3
TSEN54
SFXN3
CA12
ZNF611
CEP104
ZNF3
TBC1D3
CBX2
HMGXB4
MROH5
CTDP1

ADAMTS6
RERGL
TK2
GTPBP2
SETD3
AC093012.1
ADCY1
SMURF2
CR1
ADAMTS12
AVL9
MAGEA10
PCDH20; AL592490.1
AC004899.1
CC2D1B
FPR1
CHTOP
PTCD2
FAM83A
SMOC2
ITGA10
VNN3
PPARD
HLCS
MCF2L
ACSS3
GCSAM
SPEG
CLNK
GSR
CACNA1C
IGHMBP2
INTS7
AMPH
CCSER2
ALDH1L1
GOLGA7B
RBMY1A1
PRKN
PYCR3
FOXP2
NEIL2
SIGLEC15
HEATR5A
AC018512.1
LRRC32
MMP3
MRAS
LPAR3
DRC3
CKMT1B
PNPLA3
SHE
ANXA8L1; AC244230.1
KCNQ4
C6orf89
BBS9
UNC119B
UBN1
SLC5A7
ERCC4
MMP8
HACL1
PPP5C
IFT172
CTNNA1

CNTD1
MYO9A
OCM2
NTRK2
TMEM182
PFKFB2
C17orf49
KIAA0513
SH3BP2
SH3TC2
WDR33
GOLGB1
DCAF4
ETV3
KCNS2
APBB3
WDFY3
SLC30A8
CEP63
TNK1
ELP6
SAMHD1
PTBP3
SERTAD4
TRIM4
CDON
TREM2
MDM2
IPPK
THSD7A
EFHD2
LHX4
NBEA
EGFR
POTEC
SLC25A4
MAP3K15
TNPO1
ACOT11
SIRT5
CHD2
PURA
TBC1D3G
TRAPPC11
ERBB2
SPATA3
STAG1
UPB1
NECTIN4
IQCJ-SCHIP1
TARS2
ROPN1B
CD28
DIABLO
SOHLH2
LRP8
MCOLN3
TGFBRAP1
ZNF211
ITPKB
ENPP7
SGTA
STAT5B
KIAA1257
ADGRG2
PLEKHB2

OAS2
RPL28
SEC61G
MPP6
TNFSF8
PDE5A
SLC66A1
NRG3
C7orf61
ACSL1
ABL1
C14orf93
ZNF433
SRGAP1
UBE2H
TMPRSS7
SPATA18
CALCR
DUSP27
CRIP2
BTBD11
DIO2
MPLKIP
LRRCC1
UNC13A
SCIMP
CCT3
SLC5A10
SYNE1
KHDC4
RBMY1E
MOBP
C5orf30
NUDT16L1
PRKRIP1
GDF11
TMEM200C
RBMX2
SYT11
FBXL17
PDZD2
CEP192
RNF111
TARDBP
STAU1
ZNF160
MTO1
GRIK3
CACNB2
STXBP4
TNIP1
MAP3K4
GNG11
ITGAX
XRN1
NABP1
CANT1
CDC37
GNG7
HEATR6
RUNX2
FCRL5
SEC14L2
MBNL3
SDC3
SOX6

THRA
NAB1
SCN7A
ZFP41; AC138696.1
CCDC168
AC010642.1; AC010642.2; AC020915.1
MCF2
TRPM3
ZNF559-ZNF177
STK32B
ZNF532
ASB14
TACR3
GDPD2
SLC35C1
KIF1A
SLFN11
CYP27C1
ZDHHC21
LEKR1
FBXL16
ADAP2
TAF8
RNF139
ADAM8
RBM41
GOLGA6L1
BCL7B
RNH1
MAFG
GPR150
SHANK2
TMEM41A
PRRG3
RAPGEF5
POTEI
ZCCHC2
RBMXL2
SULF1
PRDM5
TAB3
GUCD1
ILDR2
RAB11FIP5
LPL
HCN4
LIPC
DIO1
EIF2B1
PAK4
CD84
RNF10
ZNF830
GOLGA6L22
MBP
GJC1
TANGO6
CELSR2
STAM2
AC010326.1
PDGFRA
GSG1
HELZ
PTPRJ
HOOK1
TTC14

KLHL5
KCNQ3
SPACA9
NMT1
SNX8
ZNF135
CCBE1
SARNP
DLG5
ELMOD3
C10orf105
SHOX
ATRNL1
RPAP2
ASB5
AREL1
TMEM232
G6PC
MAPKBP1
HARS2
TOMM40
AKR7L
C17orf78
HPSE
UST
LCAT
ERICH1
ST6GAL1
HMGCS1
LY9
NLRP12
ZNF570
MARS2
NFKBIA
CRTC1
AC048338.1
UPF2
GSE1
SUCLG2
FOXF2
HAUS3
PITPNC1
CX3CR1
MYLK3
LONRF1
C2orf88
CEP128
PLP1
PIK3R6
DPPA4
DENND6B
GDF7
PTCHD1
KCNJ11
TSPAN17
BRD3
BNC2
AC107021.1
WDR3
KCNJ16
CDC20B
PABPC1L2B
FABP2
LBH
SYNM
EIPR1

UGT8
ALG1
SLC25A23
GLP2R
SESTD1
NDP
FLII
ZNF486
ADAMTSL1
AP2A2
ZNF675
LINC01553
BRAF
PAX5
AAK1
AANAT
CRB1
KLK3
LTBP3
ABTB1
PASK
PHKB
ZNF776
HEPACAM
BAIAP2
ITGB5
ARSJ
KCNS3
TPD52L3
MBD1
SHMT2
VWA3B
THAP9
COL12A1
SAV1
YPEL2
RBMS1
ZNF445
ALPK2
PSD3
SAFB
NSUN4
PIGM
HECW1
TRMT2A
ZNF519
MEF2C
SENP5
SOGA3; AL096711.2
FAM174B
EIF2D
CBFA2T3
MYB
FZR1
XRCC2
ST8SIA5
DBNL
CHPT1
RPTOR
ERBB4
AMER2
LPP
CHRDL1
WNK2
RENBP
ATXN7L1

ZNF783
ITIH5
SVBP
NBEAL1
GPR107
SLC25A37
TFDP2
AP1G1
ECHDC1
SMARCC1
AMBRA1
GNAI3
COL20A1
IRF1
PTPRE
FAM20A
UNC5B
NXNL2
ZNF230
MBL2
TSPAN3
CKLF-CMTM1
SERINC3
TBC1D31
GINS3
PKD1
CHSY1
FOXR2
ARHGAP17
KCNT1
TIGD6
RARS1
C2orf91
DIAPH2
CLN8
ACVRL1
LAMP1
NFATC4
RXRA
TIMP3
ARPP21
STAG2
HRK
ACACA
ABCG1
SPAG16
ARHGEF12
CCR9
SP6
RAC2
YOD1
LNPEP
NUP93
RMC1
MRVI1
DGKH
ZNF669
CDK12
HAGHL
CACNB3
CCDC141
ZRANB3
AC022384.1
AP001781.2; ALG9
DPH3
SKP1

SEPTIN6
DRD5
TMEM30B
FYCO1
RAPH1
TBC1D5
CNTN4
CFAP57
EZH2
ATP11A
ARMC1
SYT5
BORA
SLC15A4
POU2AF1
CYP21A2
AZIN1-AS1
DRAXIN
COMMD2
MGAT1
CCDC137
SCRN2
NUTM2D
SLC26A3
SYNC
AC008060.1
PHC3
NOVA2
TMCO5A
PRKG2
TTLL1
DCAF12
NKRF
AP000769.1
PPP6R1
TNFSF15
NCF2
SH3GLB1
KBTBD6
MRPL48
FAM133B
BCL9
ZIM2
PLXNB1
SND1
F9
CNGB3
SLC25A44
CD40
LY86
ZMYND8
CCDC186
EFTUD2
HSPB11
CYP11B2
N4BP2
SPICE1
AC012314.2
ITGA1
WTIP
POU4F1
BCL2L2
SGIP1
LSP1; LSP1P1
LSG1
WDR5B

MERTK
BAG1
SLC7A14
LHX5
TMPRSS6
TP53INP2
LINC02693
BCL9L
PEX1
TUSC2
IL1RL1
RCC2
ALDH1A2
MYO16
POGK
CROT
GLP1R
TRIM37
GRSF1
RNASEK-C17orf49
CC2D1A
AC006994.2
MAGT1
EPPIN
USP54
MTF1
SCUBE1
ASAP3
LEPR
EARS2
VPS51
MAP10
AC006030.1
PEAK1
EYA1
CCNL2
GTF2F2
SIM1
CLCN2
ZFHX4
EIF2S3
AP4B1
ENDOV
BHMT2
P2RX6
CNTF
DLG4
STC1
KLHDC8B
STOM
SPOCK3
PLEKHA3
SPON1
FKBP10
RIF1
ZNF516
GGPS1
LARP1
RNF19B
MPI
LGALS3BP
NAA50
ABCC3
RHAG
ARHGEF3
SUPT3H

ADAMTS4
LDHA
PEX13
TTC5
GRIN2B
ARHGEF39
CD226
SLC38A2
SYNJ2
ATN1
SLAMF8
ORAI1
LRRIQ3
TPH1
DDX52
APAF1
TMUB2
GPR176
DPYSL5
AGK
ANKRD11
ITPR1
PABPC1L2A
NT5DC3
STX7
ATXN3
LUZP4
TMEM214
SIPA1L1
PAK3
CHCHD1
ZDHHC19
TGFBR1
WDR4
GRK2
TM4SF18
TMEM169
EBF1
MTSS2
NDUFB8; AL133352.1
TUB
ARRB1
DAPK2
SCAMP5
EREG
LRRC15
SUMF2
CENPA
CDH6
CDCA7
MAP7D2
RIMKLB
HHIPL2
CCDC127
SSTR3
KCNH6
MDM1
ZNF436-AS1
DHX37
USP53
CRYGN
MTRNR2L13
MCRIP1
ZNF630
TMEM129
G0S2

EEF1B2
SGMS1
DKC1
CCDC40
TTC7A
UBD
ATG9A
UTP23
MLLT3
TUBB3; AC092143.1
MUC4
BCL2L13
CCDC30
ZBTB39
TMEM63B
ELMOD2
GLYCTK
HARS1
UROS
NUTM2F
GTF3C2
MR1
ATG16L2
LASP1
MIEF1
IFRD1
KCTD9
TSPAN5
SLC9B2
FBXL8
HEBP2
OPALIN
GABPA
IL6
CPED1
RP13-279N23.2
USP37
AL035460.1; TMEM239
PPT2-EGFL8
IVD
VPS8
BCL7C
FP565260.3
HNRNPC
CLDN19
RNF165
ZCCHC14
DNAH2
SF1
ZMYM3
ISY1
FAAP20
LINC01588
CNOT9
ENTPD5
CNEP1R1
SLC7A2
MRPL43
NCKAP1
MAGEH1
VDR
C16orf74
BTBD6
CCDC149
GLYATL1
WDR31

SKA3
TBC1D3F
DDX19A
NEK11
P3H4
TMA16
MGAM
PPT2
ZBTB37
UHRF1BP1
RGS5
PRDM8
MUC19
FUT1
PECAM1
TSPAN15
LATS2
VPS26C
PTPN1
TYW3
PEX2
GPATCH1
BTN1A1
EZH1
VWDE
TTC31
CAPZA1
CDYL2
LRRTM4
UVRAG
MYOM2
ONECUT1
IGSF10
PCDH11Y
HGD
KLF7
GDAP2
XXbac-BCX105D18.9
COX6B1
LGMN
UBE2R2
PYGL
B3GNT4
HACD3
MTA3
DENND10
FOXO4
OPN5
PLAAT3
OTX2
SFT2D3
DSCAM
DGKI
SHLD1
AKAP1
NFIX
REG4
PLA2G5
TIMM10
NLN
BCORL1
GJA3
RREB1
PABPC1L
COL28A1
ENPP5

ZSCAN2
TOX4
FFAR4
GTPBP4
TUBGCP2
RPS15A
TAF1
IRF2BP1
MYO15A
SAMD14
TEX35
DAB2IP
KDM4C
NUP160
ERBB3
DAZL
TRIM64B
ZKSCAN8
SDK1
NAXD
CYP2C18
ATAD3C
SYK
IGFBP1
RPL3L
DHX57
AP5B1
FUT9
TBC1D1
ENTPD4
CYP27A1
FOXL2
ZNF300
LATS1
BAZ2A
SLC7A9
Z98049.1
ZNF841
FGF2
CLU
ZSCAN25
VPS28
TRAF4
CEP76
ZNF460
WNT8A
SACM1L
CX3CL1
CAPRIN2
SPTB
SNX30
C21orf62
CAV1
NAV1
RAMAC
GET1
CCL28
RETREG3
ALX4
GCFC2
TRIM38
ABCD1
TMCC1
SPARCL1
LRRC7
NCAPD2

TMEM109
WNT2B
DCP2
MRPL49
WASHC2A
GLYR1
MMUT
PPM1L
POLR1D
PSMB5
C1D
TXNDC2
LDHC
USB1
EXD3
SFTPB
CNTNAP2
NIT1
TNFRSF10B
FRK
PAX8
IRF1-AS1
TRIM71
SPECC1L-ADORA2A
SMIM14
LRRC59
LARP1B
KHSRP
CYP11B1
CAMK1D
FLRT2
KMT2A
CCDC144NL
RHBDF2
XBP1
CEBPA
FDX2
MED9
AFMID
TMEM201
SLC39A2
AKT1S1
ARL3
PHF14
NOS1AP
RPL13
SAP30BP
KCTD19
GCNT4
CLEC12A
SIRT3
HSPH1
MBLAC2
AL121758.1
H2AJ
GFAP
DPP8
NIPA1
KIAA1614
FAM83F
TLR4
KIF13B
SYNPR
SLC39A3
SUGP2
C1orf226

RP11-507M3.1
CEP126
ZNF626
HSFY1
C21orf62-AS1
C22orf34
C3orf18
IYD
CYB5RL
SUFU
DSCR4
RARG
DOCK5
DENND1C
FAAP100
CCDC15
DNAJB5
SRFBP1
TDRP
ST6GALNAC2
COQ8B
FAM162A
ANGEL2
MRPL21
TMEM213
INSR
ORAI2
WWTR1
CTU1
CRACR2A
RNF212
CBY1
MDN1
TNFAIP2
ZNF607
RGP1
PHF20L1
ST3GAL3
AC068896.3
KLF15
HLA-B
PTBP2
PPP2R2B
ZNF431
TBC1D25
JAK1
ELF5
GPD2
RBM8A
SCUBE2
VANGL1
PIGO
CHST5
NRCAM
KCNB1
LYRM4
CANX
WBSCR16; WBSCR16
DAP
ZMIZ1
EIF4A2
FRZB
CATSPER2
PLEKHA4
TSR1
CCDC85A

DPY19L4
HEMK1
GIMAP5
CAMKK2
ORM2
EGLN3
CUX1
GPX3
ZNF77
TRIM64
SLFN5
TECRL
CARF
UBR2
SHC4
GP5
OCM
STC2
COL6A1
AC253536.7
OAZ1
DSN1
GPRC5B
DIS3L
SV2C
DDR2
PPP2R5E
ASB7
AC068946.2
CNIH3
NR1I2
THSD4
KIF21B
FEZ2
KCNJ5
NXPE1
ZNF250
CRYBB2
PSEN1
GPC6
DNAJC24
AC074389.1
MCF2L2
WNK1
SPRYD3
MFHAS1
ST7
ATCAY
COA3
MFSD9
ERN1
ATG16L1
AGMAT
STK38
RFLNB
POM121
CRISPLD2
DMAC1
AOAH
CEP57
LEXM
SESN2
PLXDC1
MYO18B
RSPH9
TMEM168

CASP12
CLCF1
NUDCD1
KLHL3
CCDC112
TFF3
DPP10
PNPO
SFRP1
VAV3
SUOX
DAAM1
SCML4
ARHGAP22
MROH1
UPK3BL1
NPFFR1
ATG14
CD40LG
SMARCE1
STK17B
ZNF678
LRIG2
TMEM106B
CDKL1
CLIC6
PRODH2
UBBP4
NPTXR
HCAR1
MIGA2
GLS2
ELMOD1
SLC25A43
SMAD5
WASF2
FCHSD1
PPIL6
KRT23
SLC25A53
TNNT1
ARNT2
ZNHIT6
WDR63
RNGTT
BORCS8
APBA1
ACTN4
VPS33A
PIGX
CNTNAP1
ZNF492
SLC4A1
PRSS33
TESMIN
DDT
U2SURP
ADAM12
RBM15
ADCY7
FRAS1
SIDT1
CYCS
TECTA
ATP9B
PRKRA

KIRREL1
GYS1
DDX11
FNBP1L
PON3
ADCK1
SP140L
G2E3
NDUFS1
NWD1
PPP1R13B
CD2BP2
TOP3A
C16orf54
NDUFAB1
RAB3IP
AP3S2
SPECC1L
ZNF208
PPIAL4C; PPIAL4G
TET3
ZNF576
BBC3
RAB40B
PIKFYVE
UGT1A1; UGT1A8
PI4KB
B3GAT1
SYNGAP1
HOXA13
BET1L
FADS1
CXADR
PCYT2
TSC22D1
RHOT1
ELOVL5
SHC1
HERC2
HOOK3
CLCC1
SMO
PIK3R5
ACSBG2
AGBL4
ZIC2
SEC14L5
TMEM240
MAP3K20
OGDH
ZFP14
NECTIN2
KCNA1
CLN5
PPP3CA
CARNS1
ALDH3A2
NIBAN3
DDX53
CPLX3
GHR
FITM2
FOXJ1
CLEC4E
TMEM63C
IL17RC

C10orf120
DMTN
AC037459.1; AC037459.4
AIRE
IPO4
LMOD3
EMP2
SRSF6
TRIM41
SLCO2A1
RIN2
TMEM98
EP400
DMRT3
KBTBD3
SORT1
MFSD11
LLGL2
YWHAQ
OTUD3
YME1L1
AL357673.1
MEIS2
SLC7A3
SLC25A35
KL
IFT140
ALCAM
MYADML2
TVP23A
BCAM
CELF1
PNLDC1
ABRAXAS2
EDRF1
TMEM92
RAD18
RANBP6
FRA10AC1
PIP4K2B
GMFG
PRRC2B
LRRK2
APPBP2
LIF
OXTR
FAAP24
KIAA0355
USP19
AC007040.2
ZNF549
LIN52
ADGRG5
CLSTN2
ANXA8L2
OPRK1
PLD1
SULT1C2
WDR62
VASH1
SMIM30
NOMO1
FAM71E2
UBXN7
NGDN
HLF

MYH13
BRD8
YIPF7
ABCC11
NAALADL2
SRD5A2
SPG11
TIAM1
NOTCH2NLC
CDH24
PBX1
GOLGA8N
ZNF7
DGKD
PGM2L1
PRRX1
KLHL9
ARHGEF16
ACSF2
FAM189B
SH3D19
CDHR1
NUAK1
AL136295.4
FMN1
ASXL2
RALGPS1
TP53I13
TDRD10
SYTL4
VSTM4
SNTG2
FOXL1
CTSB
SVEP1
FANCA
C1orf198
FRMPD4
NGFR
ZGRF1
NSD1
AMMECR1
NEDD4L
KCND3
AGO2
AC008870.1
DSTYK
AP4M1
KCNG4
DNAJC16
TTC12
SELP
IFT43
CCDC169-SOHLH2
SOX10
CYB561
CHST10
DCLK2
ZNF827
TMIGD2
PPOX
SPAG9
MRPL20-AS1
HAND2
ROPN1
TCEANC

ATG7
MIP
SOD2; SOD2
ZNF627
RGS8
SH3BGRL2
TRABD2B
PEX26
LHFPL6
ADAMTSL4
SLC25A34
REXO1
MAP1A
EME2
ENTPD2
EFCAB5
FAM78A
PRDM6
RNF150
CNOT7
ITGA6
BAG3
USP35
EXOSC3
TRPM2
ZNF740
SLC17A4
RS1
NET1
TOX
CHDH
ESR1
SYT7
ZFPL1
LCA10; LCA10
DPY19L3
OLIG1
NUDCD2
AL138752.2
TCF15
FAM186B
GIGYF2
IRS2
DMXL1
SIK2
PTPN14
IBA57
ACOT9
PDE4D
POLR2J3; POLR2J3
CNTN5
GRTP1
ITGB8
GAP43
ERC2
PDE3A
PCDH11X
TMC6
XPOT
BCL6
CDR2L
CLLU1
AL159141.1
DCUN1D2
VPS53
AC140481.2; AC140481.4; PRSS40A

FRMD4A
BCL11B
PTGIR
FOXE3
ART4
EVX1
PLS3
SLC9A3R2
UCK1
RUNDC1
MAVS
ZFAND6
CD86
FAM91A1
GYG2
RAB4B
POU2F2
TSPAN14
NUDT16
TRMT11
BEND4
SNX29
FILIP1
ARGLU1
CPZ
STAT1
TBC1D10A
RBMY1D
LRRC36
FBXO30
CDCA5
GABRA1
SERINC4
DLL4
CREM
LSAMP
ZIK1
GCDH
NR2F2
BBX
TNRC6B
CYTH4
ZNF33B
ECT2
SLC25A42
MMP11
USP40
C3orf62
VWC2L
TRIP6
WDR45B
CDH3
GPR155
KLHDC8A
GSG1L
CNTROB
LYPD6B
ARHGAP20
HERC1
DOCK1
ZNF577
CCDC174
ASB11
UBR1
GPC4
VAC14

SOGA1
TMEM178B
HEBP1
SLC1A6
GFPT1
UBASH3B
ARHGAP32
EGFL8
CGNL1
AC011511.4
TMED3
HNRNPD
EXT1
ZNF528
SEC16A
NOL4
FCGRT
ZNF555
FOXJ3
UTP25
TSPAN12
COL11A2
METTL14
IL17RD
RALGAPB
PARP12
KLHDC7A
CCDC125
AC098484.3
C5orf63
GTF2H1
TAS2R5
GLRA1
HS3ST3B1
STK36
FAM102A
AC233992.2
PGF
FNIP2
DKK3
MGAT4EP
NHLRC2
ZNF430
TNRC6A
C1orf50
ZNF793
ZNF691
SEM1
LILRB3
EID1
ZC3HC1
IQCE
PTPRF
ICMT
IKBKB
MUC6
NRL
ADAMTS15
RAX2
ARHGAP1
CYP4F2
C12orf77
PRIMA1
NUTM2A
AKAP6
ACSBG1

MTMR4
CTLA4
NECAP1
BNIP1
PLA2G2D
SIRPA
IGFBP6
RELCH
TYW5
ARHGEF15
TASOR
CLEC7A
TRDMT1
GPRIN1
AC132186.1
OPCML
ALDH16A1
EYA3
BLCAP
CEMIP
SLAIN2
CEP85L
SLC35A2
CCNT1
TNKS1BP1
PLSCR1
AC211429.1; POM121C
PLEKHG4B
NOX5
CBX7
PLCZ1
ZNF589
GOLGA2
NIF3L1
JADE2
MTAP
ZFYVE28
AFDN
MAP2K7
C19orf73
PATL2
TMEM38B
SMPD3
CALN1
TMEM114
MTDH
ZNF483
C11orf45
ZCCHC17
LMBR1L
PCMTD1
GOLGA6L6
ZNF573
TBX18
RHOT2
FER
ANKK1
UTP6
COPS3
DMRT2
EXTL3
ZNF616
PI4K2B
MACC1
ARPC4
SPA17

POLE4
FGF5
GGCT
MPV17
COA7
GPX8
MIR6073
TMEM200B
SSR1
RBFOX2
NUP107
TMED10
ZNF169
C12orf45
MAFB
TMEM14B
PABPN1L
RC3H2
CCDC180
TMEM108
NACC1
PKIA
SEC62
DDX59
HOXD3
ZNF106
SEZ6L
OSBPL2
TBC1D7
SLC39A8
IL4R
ALDH1A3
RAPGEF2
EMILIN2
SMG7
SPAG8
NLRP1
AC126327.5
UNC80
FNTA
CCN1
MED24
ZNF512B
SRA1
ZNF615
MIPOL1
PPIP5K2
BRD2
DGKE
FAM53B
DNAH3
RPS6KA1
ZNF746
ATXN1
ADAM17
POLR2J2; AC105052.3
NUDCD3
RPL37A
NDRG4
GCNT3
CNDP1
S100PBP
ATP10D
MESD
HAPLN3
LZIC

CYB5D1
TRAK1
RHPN2
RABGAP1L
CD47
LAP3
TRPV3
LRRC75B
FAM104B
STX1B
ZNF620
CCDC187
VRK3
CD164
MUC13
TMEM167B
NUP205
RNF151
UBFD1
RNF8
SLCO5A1
ATP8A2
HOXC8
LARP4
SNTB1
APEX2
DEDD2
FAM228A
DDX25
KCNIP2
ZNF207
TXN2
KLF13
TOLLIP
LRRC57
IDE
NOD2
SCMH1
CERS6
PLPPR4
RP11-1407O15.2; RP11-1407O15.2
PRELP
BEND3
YARS1
POLD3
POLR3GL
TXLNB
TLCD5
LPIN1
SEMA6B
MKRN1
HACD4
PEX12
APC
CNNM2
TLR2
ZNF233
OLFM3
SBNO1
AC002451.1
B4GALT5
TRMT1L
ZBTB25
MMP19
SGK3
INO80

C8orf74
AMOT
TSHR
EPHA10
ARMCX4
AC010547.4
NRK
USP28
ATP6V0E2
CTTNBP2NL
TRAFD1
PEAR1
BACH2
HAP1
EPHB2
C9orf147
RNF123
COL27A1
NME1
ACOT8
ANKS3
CD3E
FCRL1
AC091551.1
LRTM2
CAAP1
GIGYF1
POLA2
TSPYL6
FBXL13
NOL10
EVL
SERPINA10
SYNGR1
AC016586.1
CDK3
REPIN1
NKIRAS2
YPEL3
AMER1
PDLIM5
BRWD1
ROS1
ZFAT
C1QTNF1
RUBCNL
LETMD1
TMEM54
PPIP5K1
SLC6A17
AP002990.1; MIR3654
MRNIP
FANCG
BEAN1
S1PR3
CCDC24
SLFN12L
DRD1
ACTR3C
MLXIP
KHNYN
HYAL3
ANKS1A
SLC30A9
TSPAN11
SPIB

TBCB
DHCR24
NBN
WASL
MATK
CTNND2
PTEN
COL22A1
C2orf73
ATXN7L2
TXNDC15
PKP1
KCNAB3
SYNE2
C9orf85
SLC22A4
NETO2
KLC4
CDK5RAP2
C22orf46
RPL36A
CHID1
ALPK3
FLVCR1
CTD-3138B18.4
PROM2
LMNTD1
RAB3C
CEP135
ACAD8
PLEK
LYRM2
CDH13
RC3H1
GJA1
SLC39A7
AC048338.2
BCAN
CD33
NSD2
SMPD4
FARSB
ZYG11A
TGM2
STAM
RNF141
ARHGAP30
TMEM18
NR1H3
SLC44A2
CYP2U1
TCP10
SAMD12
SNRNP70
CSGALNACT2
GASK1A
SEPTIN11
ZNF397
TTLL6
PUSL1
SYT2
SSBP4
APEH
GGA3
LPGAT1
SEC16B

SLC16A12
CASP10
MFAP3
EXOC6B
GAN
HDAC5
HLA-E
EVC2
LMLN
FRRS1L
CREG2
FAM27E3
TMEM222
SLC25A16
CSF1
MXD4
SLC8A3
DTX3L
MGAT5
MRPS11
STOML1
MTHFSD
SDR42E1
NUP153
MYH9
SEC31B
PACRGL
COPZ1
TMEM237
TMEM161B
RPS3
ADCY5
FKBP14
KLHDC7B
GPATCH2L
TSC2
NRIP1
DHX30
CPSF7
ACP2
CASC3
ZRSR2
KRAS
CATSPERG
CDX2
MTMR8
PPARG
DSG2
TMEM44
PPP2R1B
IRF6
MMAB
DCTN3
MPV17L
HCAR3
MAPT
NIBAN1
HSPA4L
SMC4
SCRG1
ZNF138
TOM1L2
ABHD15
KIAA1549L
PHOX2B
PBRM1

BICRAL
ZBTB43
IFIT2
VAPA
FAM169A
SLC34A1
FCN2
NPAP1
PARP11
CHCHD7
PAXIP1
YY1
ZC3HAV1
ZNF92
TRAPPC2L
FLRT1; AP006333.1
MCM3
KAZALD1
TENM1
DELE1
THTPA
CARD6
CCDC170
VWA5B1
SLC27A1
CBLB
MIIP
STAC2
OAS3
LIMA1
VPS39
ZNF236
CTSS
ZXDA
NEMP2
TLL2
TMTC3
LUC7L
ZNF76
ANO1
ZIC1
SOX12
ZNF286A; AC005324.4
ADH7
INKA2
ZNF587B
SMIM12
APOA5
MYO1H
LMBRD2
SYT15
SMAD7
OR11H6
PPP1R3B
SUGT1
NEU3
RGL3
CCDC27
XRCC3
LMO2
AC100839.1
GPATCH4
DUOXA1
PDPR
PHKA1
RBM20

SARM1
PDPK1
TFAP2D
ZNF792
ZFP30
PEX7
BEND7
XRRA1
TBXAS1
LGSN
ZC2HC1C
ARMC4
NICN1
NSL1
ASCC2
GFM1
FBXO17
SPRED1
TBC1D3C
MYO5A
ADAM10
GNA15
AC244102.1; GABRQ
SLC6A13
MIDN
NOX4
SZRD1
VWF
LINC00221; AC244452.1
TBX1
SEPTIN9
ZNF546
MPHOSPH8
KDM8
IQCG
MEGF9
PAPPA2
TAF1L
LAT2
NR6A1
SUGP1
HNRNPUL1
PFKFB3
PKNOX1
SIN3A
ABCB5
AHSA2P
USP2
DDA1
RNF115
TNFAIP8L1
CBX8
SLC52A3
ESD
FCAMR
UBXN2B
CSNK1D
PER2
ICOSLG
FAM118A
TRIM9
PMFBP1
TYSND1
DNAJC18
SLC24A2
SEC22B

GID8
ARHGAP19
VAV2
PTCH1
COL25A1
TFRC
DDX46
GRIA3
BHLHB9
NTRK3
ARAP3
SERPINF2
RXFP1
DCDC2
FANCF
CUX2
HYDIN
SLC10A2
RAB31
PRRC1
DIS3L2
SENP3-EIF4A1
CTCF
HEG1
CCL2
RGS9BP
ALDH3B2
FADS6
PIN4
ZBTB8A
NFAM1
ONECUT3
LMTK2
FP15737
FAM131A
TEF
FBXO43
LIMK2
HMGA2
EIF4G1
ZNF558
RANBP3
ANGPT2
B3GAT2
SLC4A10
PLIN1
ZNF618
FAM71C
RIN1
GOLGA7
SPINDOC
PHLDB1
CLOCK
MAGEC2
CEP57L1
BLOC1S1
KIFC3
CD34
WBP1L
DNAH10OS
SAP30L
BRD9
CAMKMT
LINC01555
LSM11
MSL2

ASXL3
SMYD4
ZNF701
SLC11A2
CCDC36
PLA2G2F
NPHP4
COTL1
MYD88
DLG2
TAF1A
ZC3H7B
PPP1R12B
TMEM257; TMEM257
AC118549.1
ZNF276
RRP8
AFF2
ZYG11B
RFC2
ZDHHC8
DVL3
PLXNA2
MGLL
SCFD2
SHISA6
NFASC
CLCA4
ZNF670; ZNF670-ZNF695
LONRF3
LSM6
SLC5A12
TNFSF4
COPS5
HDX
GRASP
ZFP91
SP100
PLCB1
SRXN1
LOXL2
ZNF229
EPN3
SENP2
GRIK1-AS2
MLLT6
DNMT1
EPB41L2
PCGF5
WIPF2
ZNF283
ZNF699
PCF11
FOXRED2
AK4
CDH16
NPC1L1
FMNL3
LRCH2
NAT9
MB
STXBP6
ZC3H12D
WDR25
DMBT1
SPTLC2

EXPH5
MTURN
MAP3K2
PCDH19
JMY
KMT5B
RAB3B
ADORA1
AC006455.5; AC006455.1
FGF9
ZNF322
HPCAL1
MCM9
CDH1
ZNF302
OSR1
ANKFY1
EDA
CHRNA4
SYS1
RWDD1
ZKSCAN5
LRRC74B
SSH1
ENSA
H6PD
GNPNAT1
ZNF449
CKAP2
AC006288.1
FAIM2
CTXN2
ETHE1
ZNF526
HADH
CAGE1
HNRNPM
TET2
PLEKHM1
SPIRE1
NOTCH2NLA
MRPL2
AC091167.2
RASSF2
ZNF346
DNAJC2
VPS18
TPK1
ATP8A1
ICAM1
CXCL9
SLC22A25
SIX4
RAPGEF3
PRR18
STK4
APLP2
FBXO41
PRR5L
CHRND
SRPRB
MACROH2A1
TPO
MRPS27
MPZL3
MLST8

TMEM138
MOB1A
GTDC1
U2AF1L4
VAMP3
SLC30A6
SCRN3
TRIM44
CRNN
TMEM231
SLC45A4
EID3
PRLR
JPT2
ZFYVE26
FGFR1OP
PTPRA
STOML2
INMT
GOLIM4
ALPI
LAS1L
DST
GLRX5
TTLL3
TMEM161A
PIP4K2C
ZEB2
HIF1AN
POLR2H
NCKIPSD
ABCG5
JMJD1C
SLC46A1
NR1H4
RRM1
GABRA4
SSTR1
NOTCH4
CADM1
ZNF708
PAIP2B
ARID4A
THUMPD2
APCDD1
RGS9
ZNF100
CEP41
CLVS1
YLPM1
EYS
FMO2
HAUS5
VCL
IQCJ
ZSWIM7
IRGQ
SPATA5
MAP1B
ATAT1
BRWD3
POTEH
PRPF38A
ATP6V1C2
REL
OXNAD1

CSN1S1
PDCD11
TM7SF3
CDH23
CHRD
INTS4
PEX10
KLC1
SRGAP3
AKAP13
ZNF337
DMRTC2
C6orf132
MEST
FERMT1
DYRK1A
HSPA14; HSPA14
NCAPH2
CUL2
PDS5A
DENND4C
NEDD9
PSMD13
FCER2
HDHD5
HNRNPA3
ENPP1
NOLC1
HMX3
COX11
CCND2
TKFC
CAB39L
QSOX1
UBIAD1
SLC34A2
PKDCC
CAPN10
INO80B-WBP1
SLC5A1
MYO1G
FAM114A1
FDXR
SLC9A8
ARF3
PAX3
CNP
YBX3
LMX1B
LRP4
TTC17
SLC5A2
DLGAP2
SLC16A8
CGN
SPATA13
CEP152
PDE8B
UBP1
TTPAL
MTRNR2L7
TLL1
CERS2
STPG4
WDR55
FAM9C

BLOC1S6
F12
NAA25
INTS4P2
CCDC198
CLUAP1
FAM193B
PLAC8
PTK6
LAMA4
AC010132.3; PSMA2
IGFBP4
RBMY1B
C6orf47
PDSS2
MEIOC
SNRNP27
NME3
NMD3
AL390877.1; AC115085.1; SNORA40C; AL356356.1; AL033519.1; AL031650.1; AC013429.3; AC004522.2; AL5907
PARD3B
PHOSPHO1
DNER
ETV6
HGSNAT
CD300E
PMM2
CFAP206; AL049697.1
ZNF398
ZSCAN22
ZFYVE19
PARD3
ATP2A2
MLPH
NRG1
DCAF12L1
ZNF557
ARHGAP40
AL033381.1
DGCR2
SCAF11
TAOK1
UNC45A
AC079447.1; C2orf15
LHX9
LRRD1
CNPY1
RFTN2
ANO6
NPAS3
PRKCA
EVPLL
PRDM7
CTC1
SRF
MAT2A
TUBB4A
TRPC3
SEMA4C
NDRG3
SCN5A
RUBCN
ANXA11
AC079612.1
RPS6KA3
LINC01124

| | | |
|---|---|---|
| | | PPWD1 |
| | | AC011511.1 |
| | | GPR61 |
| | | CNST |
| | | TRMU |
| | | VPS13B |
| | | SPART |
| | | SPOCD1 |
| | | DNAJC10 |
| | | PIK3IP1 |
| | | ADIPOQ |
| | | MPST |
| miRTC14 | 595 | MSRB1 |
| | | RPS11 |
| | | KCNMA1 |
| | | FAM153A |
| | | CXCL3 |
| | | COL4A5 |
| | | ITGA3 |
| | | GEN1 |
| | | CYP2D6 |
| | | FOXK2 |
| | | FOXP3 |
| | | NCOR2 |
| | | FOXE1 |
| | | USP14 |
| | | ADH5 |
| | | CDKN2A-DT |
| | | GCC2 |
| | | KLHL30 |
| | | SLC24A5 |
| | | PTP4A2 |
| | | LIMD1 |
| | | NIPSNAP3B |
| | | THRB |
| | | PWP2 |
| | | ASS1 |
| | | EMC8 |
| | | RHOH |
| | | IGSF11 |
| | | DNAH17-AS1 |
| | | PEX16 |
| | | EPHA4 |
| | | FANCC |
| | | ZNF70 |
| | | C8orf86 |
| | | CCR3 |
| | | ALDH1L2 |
| | | DNHD1 |
| | | ADRA1A |
| | | ADSS1 |
| | | FHL1 |
| | | HUS1 |
| | | HID1 |
| | | KCNK10 |
| | | RNF7 |
| | | DLGAP1 |
| | | IRF7 |
| | | XPO4 |
| | | IFNAR1 |
| | | PARD6G |
| | | CLIP2 |
| | | SAFB2 |
| | | TOR2A |
| | | CLDN10 |
| | | CYP4F12 |

CCAR2
KCNK7
KRT74
DNAJC4
APOL1
NRSN1
ZNF839
CGRRF1
TRAPPC6A
PCSK9
NDUFV2
AC145212.1
Z83844.2; Z83844.1; NOL12
GLTPD2
TMEM134
GRAMD2A
GCLC
CNOT2
RBMXL1
DDX50
CDC40
IDS; AC244197.3
ACOX3
UBE2QL1
HMBS
TBC1D16
IDO1
PPP1R15A
AP000695.1
EXOC3L4
CD276
CXCR1
FNDC9
AGRN
KRTAP10-9
PADI4
POU4F3
ZDHHC3
AFAP1
BCR
PCNP
ADAMTS8
LPIN2
RBSN
AKR1D1
SKI
UBOX5
KLF5
DDIAS
LILRA1
C16orf72
C15orf39
MTHFD1L
AMDHD1
ATXN1L
P2RX1
PDE10A
MED15
MTA1
TBC1D32
CLCN3
ESYT2
POLE
POLH
CAMTA2
AHRR; AHRR

CKAP2L
FBLN7
PPP6R3
CHIC1
MYL9
NONO
CELF6; AC009690.3
TBRG1
MAF
CSTB
PTRH2
STAMBP
SRM
NEXMIF
AKAP8
PLEKHF1
JAKMIP3
ARHGAP35
NDUFB2
WARS2
GNB1
UBE2V1
LRRC27
PRXL2B
ST3GAL5
JMJD7-PLA2G4B
UBAP2
IFRD2
STX6
MARCHF1
XKR4
PTDSS2
TBC1D23
BABAM2
HSBP1
CISD3
YIPF3
XPO7
ZBTB40
RFX2
LBR
AFF3
NBPF24
CXCL14
TUBGCP3
MTM1
HOXC13
USF2
KCNE1
PRR29
ABL2
DIP2C
SNIP1
TMEM127
NFATC1
EML1
PRKAR2A
TMEM150B
KATNAL1
UBE2I
MINK1
ZFP37
PRKCI
MRPL12
DDAH1
SKP2

SLCO3A1
OTULINL
HDDC3
AL031847.2
COMMD7
B4GALT1
KNSTRN
CLDN22
WDR75
CLDN2
KRTAP5-9
FANCD2OS
ZNF318
THAP8
ARL4C
ORMDL3
MAP11
TP73
FAM27D1
BMERB1
ABCC4
ZNF778
SRP68
NDUFS2
MRPS7
PRR14L
AP2A1
SMIM7
ATP6AP1
ADAM9
AC011484.1
EDF1
RUNDC3B
TNS1
TOM1
CCNJL
SMAD4
SLC35B3
DNAJB6
MST1
CCDC130
AKAP17A
PUS7L
OASL
SRSF8
PTBP1
PNLIPRP2
C4orf50
ZNF333
ITGAM
AC231657.1; MAGIX
EPPK1
SCYL2
PCDH1
CCDC140
NCS1
GMPPB
EAPP
NTMT1
PPAN-P2RY11
ANOS1
CD5L
IL17REL
TFDP1
SLC6A5
MOK

COX18
SPOCK2
SPEM2
CXCL12
SLC43A1
HNRNPAB
PGAP3
EFNB2
AEBP1
CDKN2AIP
MXD3
USH1G
NOS3
ARMCX6
SLC47A2
CDKN1C
ZFAND3
ADAMTS13
CDC123
MBD3
RADIL
ANKIB1
PCSK5
FMC1-LUC7L2; LUC7L2
GRM4
CNPPD1
AC120114.4
PRICKLE1
SRSF5
CASP2
PRKAG1
HLX
NRSN2
DLG1
KCNJ2
PINK1
GRAP2
WASF1
INPP5E
COX7A2L
PMPCA
RNF38
KLHL2
CSPG4
PELI2
B9D1
ZNF182
PALLD
CPLX4
TENM4
ASB13
TMEM250
RTF1
MIB2
WDR59
AC139491.7
RAB35
PDZRN3
EDNRA
ZZEF1
ATPSCKMT
ACTC1
POLDIP3
RGPD5
WDR20
IMMT

STK40
ITPRIPL2
CBARP
TCF4
NKTR
DNAAF1
MTR
NR1I3
FAM83H
RPL10
NIPAL4
KCTD10
UMPS
TMEM170A
NFRKB
BFSP2
TMEM179
RIMBP2
CACFD1
FNTB
EPHA7
GATSL2
NSMF
RAD54B
LRR1
HYPK
FDX1
WNT5A
CBX1
C17orf80
CCND1
GCNT2
SCFD1
RASGEF1A
XYLT2
SPATC1
SLC19A1
AL031666.2
LCE1C
C3orf70
ZC3H12C
TMEM53
B9D2
POLR2M
CD68
GHDC
APOF
TEAD4
PDE7B
PPP1CA
PTDSS1
SLC38A6
HEXD
MYBL1
CASQ1
INPPL1
PTTG1IP
IP6K2
MKNK2
PPP3CB
HECTD1
HDAC4
CYP2S1
CTRC
VPREB1
DNAJB7

LCN10
GCOM1
PSMG1
CEP55
RBL1; AL136172.1
NEK8
ANKRD44
SRD5A3
GUSB
CAMK4
TMEM189-UBE2V1
NMRK2
SNRNP35
SLC25A36
FBXL7
SMTN
TNNI1
MYZAP
FFAR2
PRMT9
TMEM123
GLI2
AKR1C1
CAT
ANKRD53
PTPRM
APH1B
ZC3H4
FAM118B
CFLAR
FAM210B
AC004922.1
AGPAT5
SERF2
PRKACA
PNPLA7
KPTN
CTSA
ATP8B3
AC011841.1
CLEC11A
ENGASE
ADNP2
VNN2
TMEM88
PLEKHA6
CYFIP2
ESR2
OLIG2
RPS9
SELENOT
KCNJ8
OTOG
INSIG1
ERC1
RGPD6
C8orf82
ADGRB1
UCK2
C3orf14
DIO3
CYBA
KLHDC4
DOCK2
KCTD15
AK3

PIK3CD-AS1
GIMAP8
ZNF219
MRI1
AKAP7
AC027796.3; SHPK
MANBAL
LYSMD4
BIN3
RTN4R
ST3GAL1
SRPRA
IQCH
YEATS2
TNFSF10
AC012254.2; IER3IP1
MYEF2
AGPAT3
ITSN2
MYL6
AGAP9
FGF19
LINC01465
COG2
CCDC138
YJU2
RPUSD1
MARVELD1
ACKR4
SGSM1
SH3KBP1
DNLZ
RIMS3
CADPS
ZFPM1
TWF2
SCGN
CTBP1
IRAK3
MEGF8
PPP2R2C
AGPAT4
CBY2
KCNG2
ZNF101
MAP3K10
CCNYL1
SLC5A9
IL1RAP
FOXN4
TXNDC5
OCIAD1
AVPR2
SGCG
JOSD1
TMEM187
PBK
ANKRD9
COA1
MAZ
ZNF629
ALKBH4
TLCD4
TROAP
CORO6
AC135050.2

BATF
UFC1
TRPM4
ZFR2
THSD7B
EIF4B
SLC25A25
LETM2
SORCS1
CBLN1
TCP11
CBFB
LRRK1
HOMER2
NFIB
MARCHF10
KCNAB2
REEP5
SCN2A
CCNG1
IRF5
MBD3L1
GEMIN7
PPIF
AQP12B
MICAL3
CA2
RANBP17
VXN
TMEM234
NBPF11
SLC17A2
CAB39
KLHDC10
PRKAR1B
IL16
LTO1
AC010327.2
CTBP2
PLEKHA8
USHBP1
MTSS1
CDC42BPB
UPP1
SCN8A
HSD17B8
ZNF654
ARL15
AL590822.1
XYLB
ADAMTS10
SPATA33
ZNF330
SP5
TRIT1
ZHX1-C8orf76
ETNK1
TEX30
EPS15L1
TMED1
GATAD2B
ALKBH3
CASK
SMG9
APOL3
ATP6V0E1

| | | |
|---|---|---|
| | | LRPAP1 |
| | | DUSP9 |
| | | G6PC3 |
| | | SLC17A8 |
| | | FBXO10 |
| | | DPY19L1 |
| | | LMBR1 |
| | | ST18 |
| | | RPRML |
| | | PSME3IP1 |
| | | XPO5 |
| | | SHISA5 |
| | | SLC39A6 |
| SX2 | 666 | TMEM216 |
| | | BTG2 |
| | | DECR1 |
| | | SUMO1 |
| | | CCNB1 |
| | | ERV3-1 |
| | | MET |
| | | OSBPL8 |
| | | MED13L |
| | | SCYL3 |
| | | ARL6IP1 |
| | | TCEANC2 |
| | | SERPINF1 |
| | | ATXN10 |
| | | LHFPL2 |
| | | CCL22 |
| | | CMPK2 |
| | | CDC25B |
| | | TMEM19 |
| | | TIGAR |
| | | LRGUK |
| | | EED |
| | | CD38 |
| | | FAM174A |
| | | FPGT |
| | | N4BP2L2 |
| | | CBLL1 |
| | | WDFY2 |
| | | ATP11C |
| | | PRORP |
| | | SEC23IP |
| | | CEP120 |
| | | CDC42EP3 |
| | | TLCD2 |
| | | TM6SF1 |
| | | CENPU |
| | | UBE2L6 |
| | | COMMD5 |
| | | GNB4 |
| | | OSGEP |
| | | MAD2L1 |
| | | WAPL |
| | | LMO4 |
| | | PIGL |
| | | SLC46A2 |
| | | MTG1 |
| | | TMED7 |
| | | CPVL |
| | | TRAM2 |
| | | MTMR12 |
| | | ASTN2 |
| | | GSDME |
| | | TNFAIP8L3 |

PGM1
RCAN1
HSPA2
SUMF1
RMDN3
TRIM56
CD83
KLRG1
MSS51
HMBOX1
SLC48A1
UQCRQ
SOCS4
DMBX1
ZBTB41
ARSA
PHYH
MACIR
RDH11
TRMT9B
FOXO1
RESF1
DMAC2
MKKS
STK39
FAM13A
SLC25A15
ENSG00000176593
FIG4
FAM3C
KDM7A
CYRIA
CLIC4
ITGAD
SPRY1
SERPINI1
FASTKD5
PRKD2
CHML
CFL2
BTBD1
APOOL
IPCEF1
TMEM251
PPP1R3E
IER5
GUF1
SYF2
ZBTB10
HTRA1
RAB32
IST1
RAB22A
DDX21
ARHGAP5
HDAC11
SLC9A4
METAP2
NUMB
TM9SF2
GPAM
PPP2CA
DDX17
FECH
ABHD10
SHPK

C14orf119
C2orf68
UBE2E2
CDC23
KIAA1586
SLC6A12
RASA1
CFAP97
ADAT1
MRPL51
DDO
IGSF6
SKIL
ARHGAP24
CD101
TERF1
EFR3A
VAMP7
HMGXB3
ZSCAN20
TTC33
FBXL4
RHOBTB1
RPL14
GMEB1
ARL5A
RCN2
CH25H
ZNF280D
RCBTB2
RYBP
PGBD4
PNP
HMGB1
RAP1GDS1
ATOX1
UGCG
ATRX
EIF4EBP2
MDFIC
MNAT1
RECQL
RNF170
DUSP22
STRADB
SLC39A14
EXT2
MSL3
ARL11
MAML2
LYPLA1
UAP1L1
SCOC
CTDSPL2
SHISA3
COQ7
MRPL19
NEK4
ATP6V0B
GNPDA1
MIER3
LRRC58
JAKMIP2
RNF34
FAM102B
CD36

EXO5
KIAA1143
SLC12A6
SPRTN
PAG1
PP7080
TEX261
FGD4
ZNF670
NACC2
ZNF382
NDUFAF6
STRIP2
SNRPF
NCEH1
SLC47A1
AEBP2
TRIM65
MYO10
MASTL
RUNX1
CSTF2T
STOX2
CDR2
MS4A6A
PROSER1
MFSD2A
TIFAB
FA2H
RPAP3
EFCAB14
SMARCA5
SAMD8
LNP1
MRFAP1L1
ABCE1
FAM167A
FAM229B
HSPA6
SPIN4
ATP1B2
GRHPR
TMED5
MRPL50
ANG
RIOK3
CCNT2
SPCS1
SRP19
KRI1
MRPS30
SEH1L
WDFY1
MCL1
PDIK1L
ZRANB2
HMGB2
TMEM154
MTX3
CASP3
FAM120C
ELOC
KCTD20
THAP2
PRKCE
FOSL2

FBXO21
LANCL3
GPR65
DEDD
NIFK
CCNL1
SDCBP
ORC2
TBC1D30
SLITRK5
SLC9A6
HNRNPDL
ORMDL2
IFI44L
NUP58
FOS
NEMF
ING4
ENSG00000269514
TOP2A
MRPS23
GNPDA2
CEBPG
GTF3C4
THNSL1
FAM172A
ANKRD29
SETDB2
WRN
SCML1
DOP1A
PCDH9
ZNF517
ZMYND12
FBXO4
SEPTIN14
RABL3
HERC4
RPL41
ABRAXAS1
PAICS
MYCBP
RNF125
RRM2B
HSD17B12
NDUFA9
RRAGD
VOPP1
FCRLA
SAMD9
HDGF
ARID4B
GMDS
FICD
DLAT
HIP1
TRPS1
KPNA6
SERBP1
TNFSF14
VTA1
FBXO32
ITPRIP
IWS1
TDP2
ALDH9A1

ADD2
C11orf54
SAR1B
DPCD
SGPP1
FAM161B
WDSUB1
CNBP
G3BP1
NUP155
UBE2G1
ARRDC2
PAPOLA
TTC37
HLA-DRB5
RWDD2A
SDAD1
PRR16
ZNF529
ATP5MC1
PLEKHO1
BBS7
SERPINB9
GLE1
GUCY1A1
POLR1E
RNF217
ZNF25
HNRNPR
ZNF605
UBXN4
PACC1
RGL1
ATG10
PRXL2C
AFAP1L1
EZR
ARIH1
CRPPA
KIF2A
OCRL
PRRG1
FXN
AP3B1
UBL7
FAM126B
CBX6
PHACTR1
BCAP29
HEXIM1
C6orf62
ZNF891
MRPS28
C12orf65
ZNF417
FAM114A2
GLDN
NCAPD3
BRCC3
YPEL5
RBM26
ANKRD22
NAPB
SLC39A9
GPATCH11
CCDC90B

UBE2D3
GALNT2
REPS2
PRKDC
SLC30A7
PLEKHO2
MRPS9
FAM107B
ADGRE2
EPSTI1
FASTKD2
ZNF354A
HIPK3
TXNL4A
LIMS1
ABRACL
ABHD18
ZBED6
RSPH3
PFN4
TPST2
RPL24
YES1
FAM120AOS
SDHAF1
RCC1
RGS18
ZDBF2
ELL3
CHMP4C
METAP1
WEE1
SPAST
RGS1
RERE
TIMM8A
SRRM1
HACD1
CDC27
SIKE1
KDM5B
KPNA4
RABIF
ASL
TFAM
PIGH
ENSG00000272195
HNRNPA2B1
TMEM70
MARCHF5
STIM2
MS4A4A
SPTY2D1
SPOP
BLVRA
ZNF268
MED7
DOK1
SLC25A17
SRD5A1
RIOK2
ABHD13
LRRC37B
WDR77
KCTD11
HDAC8

STARD4
DNAJC27
CLEC17A
RALGPS2
PMPCB
HYLS1
FBXW7
LRRC8B
RPA3
ZNF396
ESCO2
SP3
GAB3
PET117
GLS
SPAG1
PPP6C
TMEM115
PLPP3
PLGRKT
SDE2
UBE2D4
NCOA2
BNIP3L
LCP1
USP12
PREPL
RAB39A
PPIC
MXD1
AZI2
COQ4
MMADHC
COL8A2
JDP2
ERRFI1
SUCNR1
COMMD9
GALNT1
SNAPC3
HAVCR2
LACTB
HACD2
MSANTD4
EDEM3
SLC16A1
SCO1
ZNF440
AP1AR
TMEM63A
MYADM
CCDC59
EHF
GTF2A1
SLC28A3
ROGDI
HMGB3
NDUFAF3
ARL1
EXOC5
RAB8B
ZNF506
NOM1
ASCC3
HPS6
APOL6

FBN2
KIAA1958
MIS18A
P3H1
KLHL23
ARMCX5
AMN1
ZNF141
SRI
LDLRAD4
TFCP2L1
SCIN
ATP13A3
CSDE1
RNF215
SPOPL
TRMT10B
ODR4
JAG1
HSPA4
VPS41
C11orf58
C1orf21
ZW10
RNF20
USP9X
NEK7
SLBP
UBL4A
STK38L
RORB
SLC9A7
AKIP1
ANKRD46
SNX4
TIMM10B
BZW1
DDX60
CENPS
ZCCHC7
STXBP5
ZNF124
PRKX
CCSAP
STING1
ZBTB26
GALM
ACO1
TULP4
OSTM1
TXNL1
CISD2
PNRC1
MAT2B
MTBP
NEDD1
BCOR
PNPLA8
RP2
SNTB2
FUT10
PDCD10
SMYD2
CHMP4B
GALNT4
MED23

EEF1A1
MTPAP
ACSM5
TMSB4Y
GCNA
KATNBL1
CPNE8
ZC3H13
ZNF45
ARPC2
TMBIM4
OGT
PHIP
NIP7
SESN3
DAB2
ZNF597
CSTF2
UGDH
KNL1
DENR
BRIP1
UQCC2
GOLGA8H
GSTK1
CEP97
ZNF844
ITPA
SRSF1
CIAO2A
MTRF1L
BCL2L1
HECTD3
FGF18
CHIC2
ZNF367
DNAJC28
ATP5PB
GTPBP10
DNAJB9
PAFAH1B2
TIMM8B
TMEM170B
MINDY2
NDUFAF5
C1orf35
ZCCHC8
HFE
AKAP5
IFT81
MLEC
OAF
RNF121
TMEM30A
SYT17
CCDC171
SNRPD3
RNF149
RAD21
TAPT1
TMEM199
ZNF770
FANCE
GMPS
PTGS1
TACO1

ATE1
DCTN5
ING1
BCAT1
C18orf54
MALSU1
THAP6
ANXA4
NANOS1
UAP1
IRAK4
KCNJ15
FCHO2
KAT6B
VHL
UBXN2A
DCLRE1C
TTC30A
BNIP5
